# Inferring causal pathways among three or more variables from steady-state correlations in a homeostatic system

**DOI:** 10.1101/278101

**Authors:** Suraj Chawala, Anagha Pund, B. Vibishan, Shubhankar Kulkarni, Manawa Diwekar-Joshi, Milind Watve

## Abstract

Cross-sectional correlations between two variables have limited implications for causality. We show here that in a homeostatic system with three or more inter-correlated variables, it is possible to make causal inferences from steady-state data. Every putative pathway between three variables makes a set of differential predictions that can be tested with steady state data. For example, among 3 variables, A, B and C, the coefficient of determination, 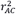 is predicted by the product of 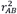 and 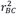 for some pathways, but not for others. Residuals from a regression line are independent of residuals from another regression for some pathways, but positively or negatively correlated for certain other pathways. Different pathways therefore have different prediction signatures, which can be used to accept or reject plausible pathways. We apply these principles to test the classical pathway leading to a hyperinsulinemic normoglycemic insulin-resistant, or pre-diabetic state using four different sets of epidemiological data. Currently, a set of indices called HOMA-IR and HOMA-β are used to represent insulin resistance and glucose-stimulated insulin response by β cells respectively. Our analysis shows that if we assume the HOMA indices to be faithful indicators, the classical pathway must in turn, be rejected. Among the populations sampled, the classical pathway and faithfulness of the HOMA indices cannot be simultaneously true. The principles and tools described here can find wide application in inferring plausible regulatory mechanisms in homeostatic systems based on epidemiological data.

## Introduction

In the field of biomedicine, the nature of causality, and the use of correlations as an evidence for causality are much debated (1–6). There have been many attempts to develop sound methods to address questions of causal inference from correlational data which include Hill criteria (7), path analysis (8–11) the use of instrumental variables (12), Granger causality (13), Rubin causal model (14), or additive noise models (15). Hill criteria are a set of common sense criteria useful to avoid making misguided inferences. Path analysis generally assumes a direction of causality, and is useful in determining the contributions of different causal pathways to a process or a resultant variable. It generally assumes directed acyclic paths and its application to pathways with loops and cycles is difficult. Methods like Granger causality depend upon the assumption that the cause always precedes effect and that the variables show some degree of chaos or turbulence, so that there are notable events like sudden peaks in the variables, which can be tracked using longitudinal data. In evolved systems in which predictive adaptive responses are possible, the assumption that cause always precedes effect is questionable. Another class of methods like Propensity Score matching based on the Rubin causal model works well to estimate the effect of a causal factor, but does not take into account unobserved factors. The Rubin Causal Model also incorporates the structural equations model as it includes non-parametric forms as well (16,17). Models like additive noise can suggest the direction of the arrow of causality between two variables, but they require the assumption that either A causes B, or B causes A, without any confounding, looping or circularity (18,19).

More specifically, here we look at homeostatic systems which are extremely common in fields such as physiology. Homeostatic systems have a unique problem for causal inferences. Causal inference can be based on time-series analysis with longitudinal data (19,20). Longitudinal data are of little use however, if the time taken to reach equilibrium is smaller than the observational window, or if the system is already in a steady-state. Most homeostatic systems have negative feedback or some loop structures, because of which methods assuming acyclic causal paths or freedom from confounding are not applicable.

Although the use of correlations to infer causality is doubted, intervention experiments are generally taken as a convincing evidence of causality. However, causality in steady-state can be substantially different than causality in a perturbed-state and inferences from a perturbation experiment may not be applicable to steady-state causality. This necessitates a set of tools to infer causality in a steady state which is independent of perturbing interventions. We argue in this paper that it is possible to infer causal relationships among three or more variables from cross-sectional data in a homeostatic system in which the variables and their relationships are stable in time.

## Motivation

Our motivation and the need for this tool came from some debated causal pathways in the pathophysiology of type 2 diabetes (T2D). According to the classical view, obesity-induced insulin resistance is primary, and rise in insulin levels is a compensatory response to insulin resistance, mediated by raised levels of glucose (21,22). This is contested (23), with increasing evidence suggesting that hyperinsulinemia precedes insulin resistance (24–28).Therefore the causal pathways between insulin levels, insulin resistance and plasma glucose are uncertain. There is also evidence of neuronal signals affecting insulin production on the one hand, and controlling glucose production by the liver partly independent of insulin on the other. Therefore, the causal relationship between insulin resistance, hyperinsulinemia and hyperglycemia needs to be re-examined (reviewed by (23)).

Elucidation of the causal pathway for a pre-diabetic or diabetic state is critical at the clinical level because the current approaches to medication are designed assuming one pathway but have largely failed to cure diabetes. If it is possible to determine causality reliably, it can potentially change diabetes medicine. It has long been recognized that levels of glucose and insulin are under homeostatic control, and that fasting is a steady state (29–32). With substantial data available on fasting levels of glucose and insulin from different populations, along with many other variables, a tool for inferring causality from a set of inter-correlated steady-state variables would help understand, and thereby better control T2D.

Beyond the specific problem of causality in pre-diabetes, a set of methods that can infer causality from steady-state data will find a large number of applications, not only in physiology and disease, but in many other areas of science. Although our investigations began with the pathophysiology of diabetes, the emerging principles are generalizable and valuable for inferential statistics in general.

## Methods

We show here that when three inter-correlated variables are considered together with two or more causal arrows connecting them to make a causal pathway, each of the possible pathways makes a set of differential predictions by which the pathways can be differentiated from each other. Our approach to develop a method of inferring causality from cross sectional regression correlation parameters comprises following steps.

1. We first list the perceived possible hypothetical causal pathways among three variables.
2. For each pathway, we write a set of causal equations arising out of the hypothetical pathways. Steady-state solutions of these equations lead to a set of four general, and a few pathway-specific predictions. Each pathway therefore has a unique combination of such predictions or a prediction signature by which it can be differentiated from other pathways.
3. We test, using simulated data generated from assumed causal pathways, the conditions under which the predictions can be used to accept or reject a pathway reliably.
4. Based on these results, we suggest ways of handling multivariate data and infer causal networks among them.
5. We apply this logic to the specific case of pre-diabetes to examine the pathway classically thought to give rise to this condition.

### Baseline assumptions and nomenclature

We consider three variables labelled A, B and C. Additional variables if needed to describe a pathway will be labelled X, Y and so on. All causal relationships represented by a single arrow are assumed to be linear, and all primary input variables are assumed to be normally distributed. In a given operation, the slopes of causal pathways are assumed to be constant; the errors in causal pathways are assumed to be distributed normally, with a mean zero and a constant standard deviation, and no covariance with each other. We assume that the errors are caused by variation in individual responses, and that a given individual’s response is consistent in time sufficiently long to reach a steady state. So the errors are randomized over the population, but for a given individual, they are constant in time. We assume no measurement errors in the baseline models. Since all our predictions are related to correlation coefficients and regression slopes, we will ignore the intercepts for the sake of simplicity in deriving many of the predictions.

### The possible pathways

A variety of cyclic and acyclic pathways can exist in three variables. Fig 1 shows the simple primary pathways that can exist. More can certainly be constructed by combinations of the primary pathways. It is also possible to consider permutations of the three variables. For example, the linear pathway among three variables can itself be written in six different ways. Here we restrict to the primary pathways assuming a fixed sequence of the three variables denoted by A, B and C. The principles that we derive from this set of primary pathways can be extended to more complex pathways.

**Fig 1.**
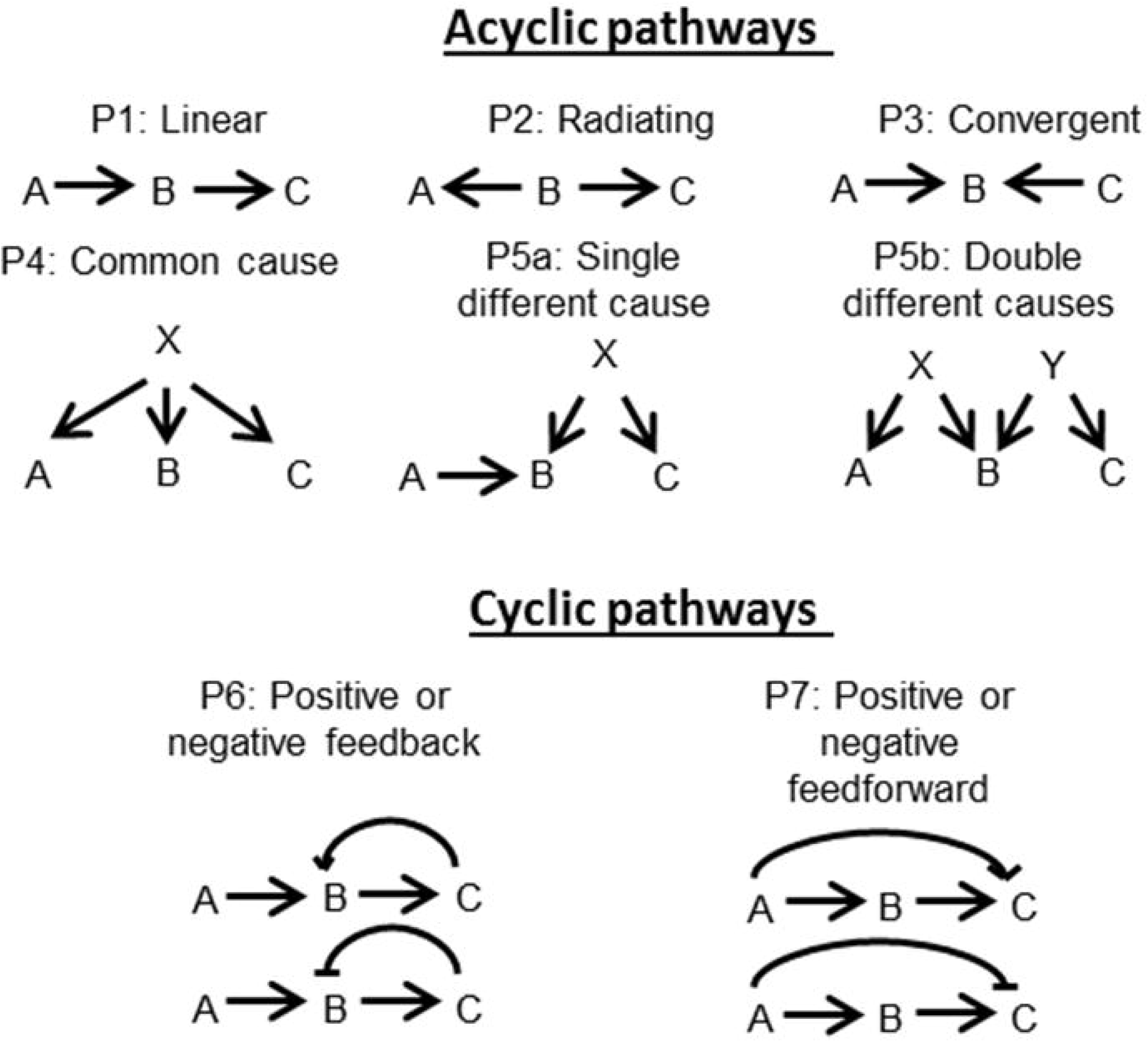
Possible primary causal pathways between three variables. More complex pathways can be visualized by combinations of the primary ones.

### Causal equations versus regression equations

Based on hypothesized pathways, we can write specific causal equations for each. The causal equations are derived from the hypothesized pathway, while the regression equations can be obtained from the given cross-sectional data using regression and correlation analysis. Our causal equations are similar to the structural equations of (17). However, they differ in their interpretation and treatment. In structural equations, the left hand terms are effects and right hand terms are causes, and the two cannot be algebraically transferred without changing causal interpretations. In our approach, after finding equilibrium solutions, we can carry out algebraic operations freely in order to obtain testable predictions. The parameters of the regression equation are not necessarily identical to those of the causal equations (Table 1). For example, for a hypothesized pathway Y = mX + C, m is the causal slope, while the regression slope would be underestimated if there is post-effect variability in X (33). Such a bias in the slope is important in making and testing predictions. In the following section, we show that the parameters of causal equations hold pathway-specific relationships with the parameters of regression equations based on which, pathway-specific predictions about the regression correlation parameters can be made.

**Table 1.**
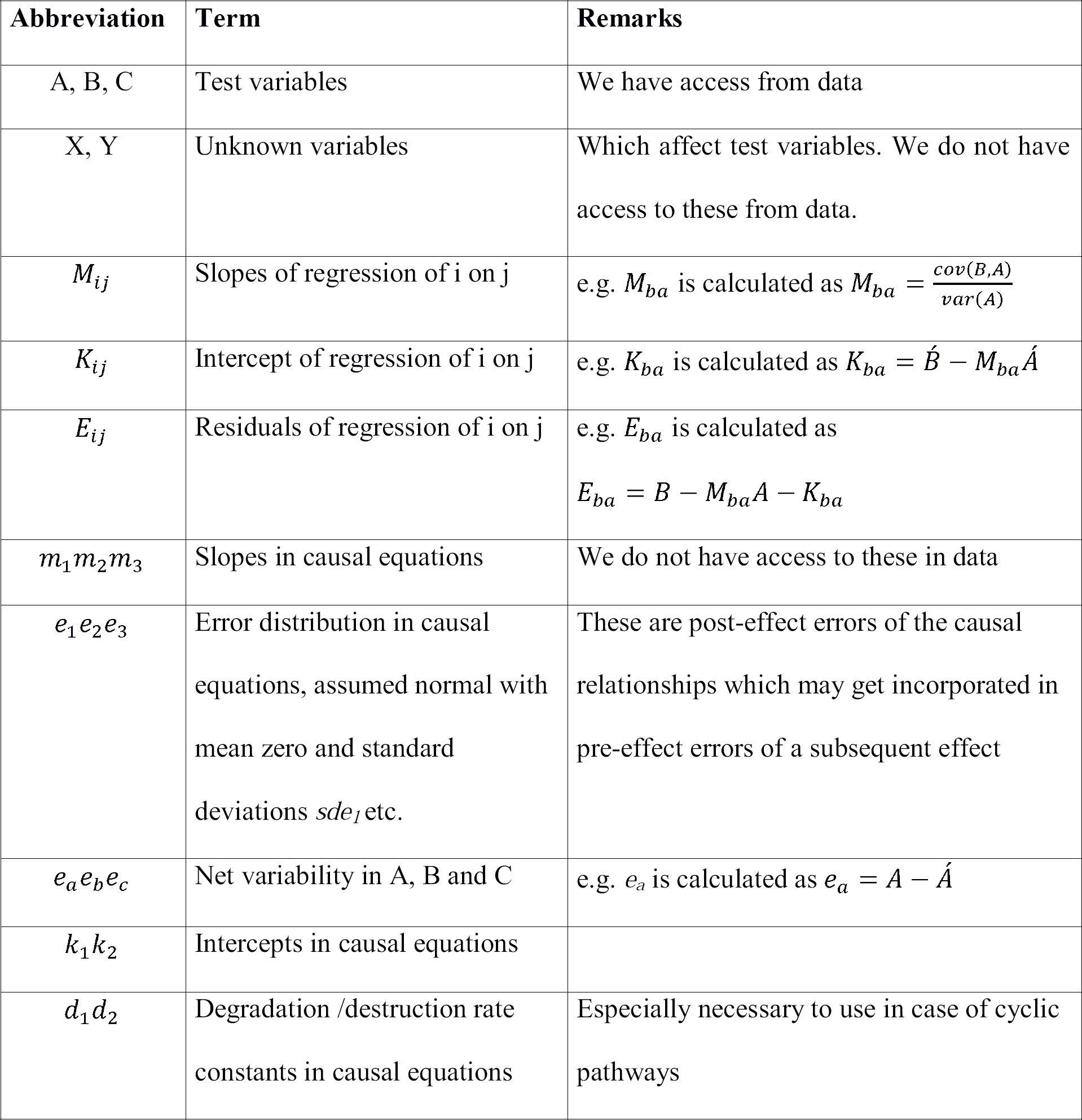
List of abbreviations used.

| Abbreviation | Term | Remarks |
| --- | --- | --- |
| A, B, C | Test variables | We have access from data |
| X, Y | Unknown variables | Which affect test variables. We do not have access to these from data. |
| $M_{ij}$ | Slopes of regression of i on j | e.g. $M_{ba}$ is calculated as $M_{ba} = \frac{cov(B,A)}{var(A)}$ |
| $K_{ij}$ | Intercept of regression of i on j | e.g. $K_{ba}$ is calculated as $K_{ba} = \hat{B} - M_{ba}\hat{A}$ |
| $E_{ij}$ | Residuals of regression of i on j | e.g. $E_{ba}$ is calculated as<br>$E_{ba} = B - M_{ba}A - K_{ba}$ |
| $m_1m_2m_3$ | Slopes in causal equations | We do not have access to these in data |
| $e_1e_2e_3$ | Error distribution in causal equations, assumed normal with mean zero and standard deviations $sde_1$ etc. | These are post-effect errors of the causal relationships which may get incorporated in pre-effect errors of a subsequent effect |
| $e_ae_be_c$ | Net variability in A, B and C | e.g. $e_a$ is calculated as $e_a = A - \hat{A}$ |
| $k_1k_2$ | Intercepts in causal equations | |
| $d_1d_2$ | Degradation /destruction rate constants in causal equations | Especially necessary to use in case of cyclic pathways |

Table 1 legend: Parameters of causal equations are denoted by small letters and those of regression equations by capital letters.

For ensuring steady-state, we assume that a given variable has a rate of formation/increase and a rate of degradation/decrease. If the rate of degradation is positively dependent, or the rate of formation negatively dependent on the standing level, then the variable invariably reaches a steady state determined by the set of input parameters. Such steady states are characteristic of homeostatic systems, and this principle is central to our methods.

For example, in a linear pathway 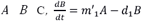 and 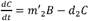. At a steady-state the net change in any variable is zero. Therefore, the steady-state levels of *B* and *C* will be 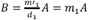 and 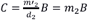 respectively.

In simple cases, we need not explicitly include the rates of degradation in the equations but directly use parameters *m*_*1*_ and *m*_*2*_. For pathways involving loops and feedbacks the relationships between variables are more complex and for such cases we will explicitly use the degradation constants in the causal equations for ensuring steady-states.

### Making predictions from steady-state solutions

We make four general predictions across all pathways and then formulate a null hypothesis for each. In addition, there are certain pathway specific predictions that will be discussed along with the description of the corresponding pathway. The four general predictions are:

1. Whether 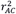 can be estimated from the product of 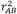 and 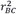.
2. Whether slope *M*_*ca*_ can be estimated from the product of the slopes *M*_*ba*_ and *M*_*cb*_.
3. Whether the residuals of the regression of *B* on *A* (*E*_*ba*_) are correlated with those of *C* on *B* (*E*_*cb*_): The errors or residuals in a regression are assumed to be random independent errors. However, we will show below that if there are loops, convergent or confounding elements in a pathway, *E*_*ba*_ and *E*_*cb*_ do not remain independent. Based on the nature of dependence between *E*_*ba*_ and *E*_*cb*_, presence of, and possible nature of the loops and convergence can be inferred.
4. a. Whether correction for *A* improves or reduces the correlation of *B* with *C*, i.e. whether 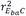 is greater or lesser than 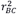. b. Whether the extent to which 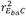 is greater or lesser than 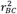 can be predicted by 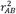. We will now state how each of the pathways makes specific predictions. For detailed formal proofs and derivations refer to S1 Text.

### Making and testing analytical predictions

#### Acyclic pathways

#### Linear Pathway (P1)

The causal equations for a linear pathway are:

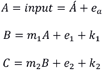

Where *e*_*a*_, *e*_1_, *e*_2_ are not correlated.

Regression parameters can be derived from the causal equations as follows. Since in regression of *B* on *A*, the slope = *cov (A, B)/var A*,

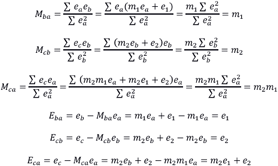

For linear equations, there is little difference between the causal equations and regression equations (Table 2). The regression equations therefore become

**Table 2.** Relationship between the causal and regression equations for linear pathway.

| Slopes | Errors |
| --- | --- |
| $M_{ba} = m_1$ | $E_{ba} = e_1$ |
| $M_{cb} = m_2$ | $E_{cb} = e_2$ |
| $M_{ca} = m_1 \cdot m_2$ | $E_{ca} = m_2e_1 + e_2$ |

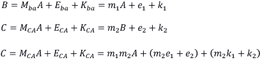

Prediction R1: Based on the equations above and Table 2, it can be shown that *r*_*AC*_ – *r*_*AB*_*r*_*BC*_ *=0* (See S1 Text ‘Linear pathway: Prediction R1: Proof 1’ for formal proof). Prediction R2: From Table 2, it is obvious that the slope *M*_*ca*_ can be predicted from the product *M*_*cb*_*M*_*ba*_ ; *M*_*ca*_ *− M*_*cb*_*M*_*ba*_ *= m*_*1*_*m*_*2*_ *− m*_*2*_*m*_*1*_ *= 0*.

Prediction R3: From Table 2, as there is no covariance between *e*_1_ and *e*_2_,

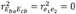

Prediction R4: For a linear pathway, it can be shown that

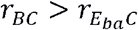 and further, 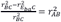

#### Radiating pathway (P2)

The causal equations for this model would be

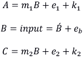

Note that the relationship between causal parameters and regression parameters is substantially different in this pathway than the linear pathway (Table 3). For example, the causal slope is 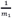, but there is an underestimation of the slope during regression which is predicted exactly by 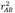.

**Table 3.** Relationship between the causal and regression equations for radiating pathway.

| Slopes | Errors |
| --- | --- |
| $M_{ba} = \frac{1}{m_1} r_{AB}^2$ | $E_{ba} = e_b(1 - r_{AB}^2) - \frac{1}{m_1} r_{AB}^2 e_1$ |
| $M_{cb} = m_2$ | $E_{cb} = e_2$ |
| $M_{ca} = \frac{m_2}{m_1} r_{AB}^2$ | $E_{ca} = m_2 e_b(1 - r_{AB}^2) - \frac{m_2}{m_1} r_{AB}^2 e_1 + e_2$ |

However, this difference is not detectable from cross-sectional data alone. Therefore, the standard four testable predictions of this pathway remain similar to the linear pathway. We will describe later that differentiating between pathways P1 and P2 is possible using a different strategy.

Prediction R1: From Table 3, it can be shown that_Ac_ − r_AB_.r_Bc_ = 0.

Prediction R2: From Table 3; we see slope M_ca_ can be predicted from the product M_cb_M_ba_

Prediction R3: From Table 3, 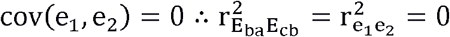

Prediction R4: As formally shown in S1 Text (‘Radiating pathway: Prediction R4: Proof 2’), 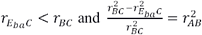

#### Convergent pathway (P3)

The causal equations for this model would be

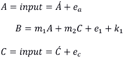

where *e*_*a*_, *e*_1_, and *e*_*c*_ are uncorrelated.

Regression parameters derived from the causal equations are given in Table 4.

**Table 4.** Relationship between the causal and regression equations for Convergent pathway.

| Slopes | Errors |
| --- | --- |
| $M_{ba} = m_1$ | $E_{ba} = e_1 + m_2 e_c$ |
| $M_{cb} = \frac{1}{m_2} r_{BC}^2$ | $E_{cb} = e_c (1 - r_{BC}^2) - \frac{1}{m_2} r_{BC}^2 (m_1 e_a + e_1)$ |
| $M_{ca} = 0$ | $E_{ca} = e_c$ |

There are two pathway specific predictions for the convergent pathway, shared only by the different cause pathway. Firstly, we expect no correlation between *A* and *C* from this pathway, unless there are additional external pathways linking the two. The other unique feature of this pathway is that both *A* and *C* have independent causal influence on *B*. As a result, the effect of *A* adds to the error in the correlation between *B* and *C* and similarly, the effect of *C* contributes to the error in the correlation between *A* and *B*. As a result, 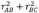 cannot be greater than 1, as shown below:

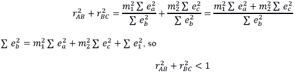

This prediction is so robust that if 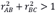, the convergent pathway can be rejected right away. Since we assume *A* and *C* to be independent input variables we assume no correlation between them. However, if they are correlated due to some cause other than this pathway, only then 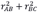 can be greater than 1.

Prediction R1: Unlike pathways P1 and P2, for the convergent pathway, it can be seen that

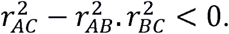

Prediction R2: Since the expected slope *M*_*ca*_ is zero, and both *M*_*ba*_ and *M*_*cb*_ are non-zero, their product is not a predictor of *M*_*ca.*_

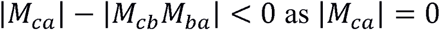

Prediction R3: The correlation 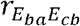 is predicted to have the same sign as M_cb._

Prediction R4: It can be shown that (a) 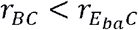 and further, (b) 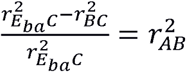.

Because this expression differs from R4 (b) of the earlier pathways, we can use a more generalized form for R4 (b) as 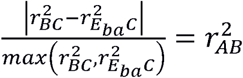.

#### Common cause pathway (P4)

The causal equations for this model would be

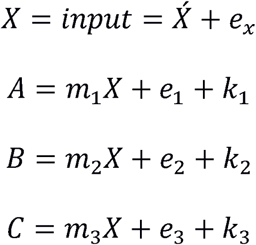

where *e*_x_*e*_1_*e*_2_*e*_3_ are not correlated.

It needs to be noted that *e*_1_*e*_2_*e*_3_ are important in defining this pathway. If *e*_2_ is negligible the pathway approximates to the radiating pathway with *B* being the mediator between *A* and *C*. Similarly, at small *e*_1_, *A* becomes the mediator and at small *e*_*3*_, *C* becomes the mediator in a radiating pathway. For the way we have defined our predictions, *e*_2_ is the most important error term in this pathway (Table 5).

**Table 5.** Relationship between the causal and regression equations for common cause pathway.

| Slopes | Errors |
| --- | --- |
| $M_{ba} = \frac{m_2}{m_1} r_{AX}^2$ | $E_{ba} = m_2 e_x (1 - r_{AX}^2) + e_2 - r_{AX}^2 \frac{m_2}{m_1} e_1$ |
| $M_{cb} = \frac{m_3}{m_2} r_{BX}^2$ | $E_{cb} = m_3 e_x (1 - r_{BX}^2) + e_3 - r_{BX}^2 \frac{m_3}{m_2} e_2$ |
| $M_{ca} = \frac{m_3}{m_1} r_{AX}^2$ | $E_{ca} = m_3 e_x (1 - r_{AX}^2) + e_3 - r_{AX}^2 \frac{m_3}{m_1} e_1$ |

One very special feature of this pathway is that qualitatively it is highly symmetric with respect to all the three variables *A, B* and *C*. This means that any permutation of them does not change the qualitative nature of any prediction. This can be used as a pathway specific prediction and a distinct signature for this pathway.

Prediction R1: It can be shown that 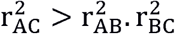 for this pathway.

Prediction R2: |M_ca_| − M_cb_M_ba_ v0

Prediction R3: The sign of the correlation 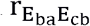 is decided by the signs of m_1_ and m_2_. When both have the same signs 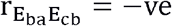 and when they have opposing signs 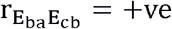. In other words the correlation multiplied by the sign of M_ca_ is always negative.

Prediction R4: For this pathway (a) 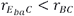 and (b) 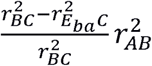

#### Single different cause pathway (P5a)

The causal equations for this model would be

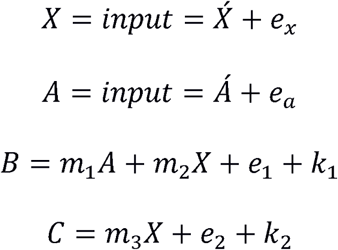

where *e*_x_*e*_1_*e*_2_*e*_3_ are not correlated. Regression parameters derived are in Table 6.

**Table 6.** Relationship between the causal and regression equations for Single Different Cause pathway.

| Slopes | Errors |
| --- | --- |
| $M_{ba} = m_1$ | $E_{ba} = m_2 e_x + e_1$ |
| $M_{cb} = \frac{m_3}{m_2} r_{BX}^2$ | $E_{cb} = m_3 e_x + e_2 - m_3 r_{BX}^2 \left( \frac{m_1 e_a}{m_2} + e_x + \frac{e_1}{m_2} \right)$ |
| $M_{ca} = 0$ | $E_{ca} = m_3e_x + e_2 - m_1e_a$ |

Two specific predictions of this pathway shared only by the convergent pathway (P3) are that

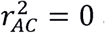 and that 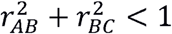.

Prediction R1: 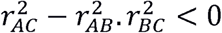

Prediction R2: |*M*_*ca*_| < *M*_*cb*_*M*_*ba*_ V

Prediction R3: The correlation 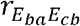 is predicted to have the same sign as M_cb_.

Prediction R4: (a) 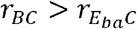 and further (b) 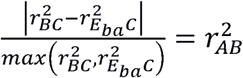 is true.

#### Double different causes pathway (P5b)

The causal equations for this model would be

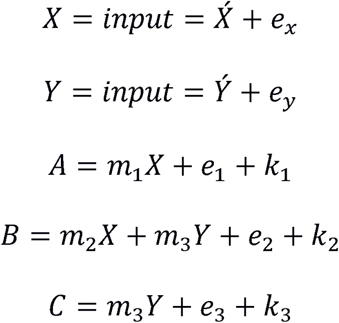

where *e*_x_*e*_y_*e*_1_*e*_2_*e*_3_ are not correlated. Regression parameters derived are in Table 7.

**Table 7.** Relationship between the causal and regression equations for Double Different Cause pathway.

| Slopes | Errors |
| --- | --- |
| $M_{ba} = \frac{m_2}{m_1} r_{AX}^2$ | $E_{ba} = m_2 e_x (1 - r_{AX}^2) + m_3 e_y + e_2 - \frac{m_2}{m_1} r_{AX}^2 e_1$ |
| $M_{cb} = \frac{m_4}{m_3} r_{BY}^2$ | $E_{cb} = m_4 e_y (1 - r_{BY}^2) + e_3 - \frac{m_4}{m_3} r_{BY}^2 (m_2 e_x + e_2)$ |
| $M_{ca} = 0$ | $E_{ca} = m_4 e_y + e_3$ |

Two specific predictions of this pathway shared only by the convergent pathway (P3) are that 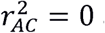 and that 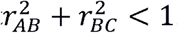.

Prediction R1: 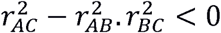

Prediction R2: |*M*_*ca*_| < *M*_*cb*_*M*_*ba*_ v

Prediction R3: The correlation 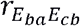 is predicted to have the same sign as M_cb_.

Prediction R4: 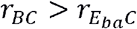 and further 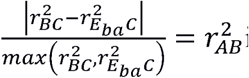 is true.

It can be seen that all predictions of pathways P5a and P5b are identical and henceforth we will treat both of them in a single group as pathway P5.

### Pathways with loops

In pathways with loops, since there is a cyclic dependence between the variables we begin with differential equations with variable-specific constant rates of disintegration that assure steady states. This set of equations is then used to derive equilibrium solutions.

#### Positive or negative feedback pathway (P6)

The causal equations for this model would be

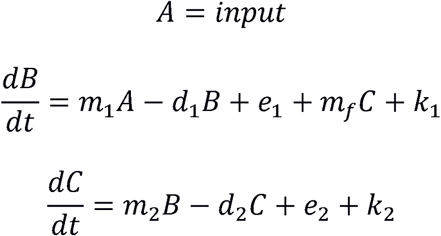

where *e*_a_*e*_1_*e*_2_ are not correlated, and *d*_1_ and *d*_2_ are positive. For a consistent definition of feedback, we assume *m*_1_ and *m*_2_ to always be positive, and that the sign of *m*_*f*_ decides whether it is a positive or negative feedback loop. Feedback loops depend crucially on the relative strength of the forward versus backward causation. If the feedback term, i.e. the effect of *C* on *B* is weak, it approximates to a linear pathway, and if the forward term i.e. effect of *B* on *C* is weak, it approximates to a convergent pathway. Therefore the predictions of linear or convergent pathways can be expected if the forward or feedback links respectively are weak. Additionally however, the negative feedback pathway is associated with a problem of definition. If the feedback effect of *C* on *B* is stronger than the effect of *B* on *C,* the signs of the slope in the causal and regression equations could be opposite, implying that while *m*_2_ is positive, *M*_*cb*_ could become negative. This happens when

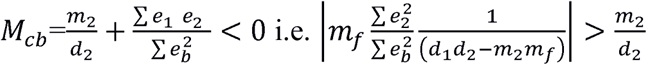

This results in a paradoxical transformation of a causally negative feedback into an apparent positive feedback since the sign of the slope and that of the feedback effect is the same. Further, when the negative feedback is much stronger than the forward effect, the predictions of convergent pathway are more applicable than the predictions of negative feedback pathway. At equilibrium where both 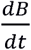 and 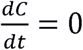, the equilibrium concentrations of *B* and *C* are given by

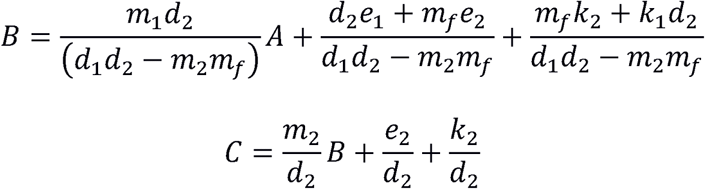

For simplification we take 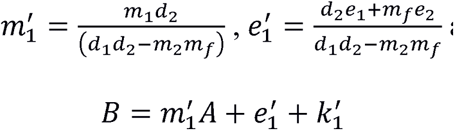 and 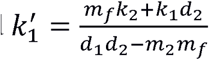

Similarly 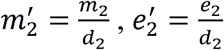 and 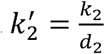

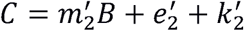

It should be noted that 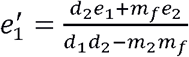 and 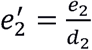 share *e*_2_, and would therefore co-vary. The sign of this covariance is decided by the sign of *m*_*f*_, i.e. whether the feedback is positive or negative.

Regression parameters can be derived from the above as in Table 8.

**Table 8.** Relationship between the causal and regression equations for Positive or negative feedback pathway.

| Slopes | Errors |
| --- | --- |
| $M_{ba} = m_1 = \frac{m_1 d_2}{(d_1 d_2 - m_2 m_f)}$ | $E_{ba} = e_1 = \frac{d_2 e_1 + m_f e_2}{d_1 d_2 - m_2 m_f}$ |
| $M_{cb} = m_2 + \frac{\sum e_1 e_2}{\sum e_b^2}$<br>$= \frac{m_2}{d_2} + \frac{\sum e_1 e_2}{\sum e_b^2}$ | $E_{cb} = e_2 - \frac{\sum e_1 e_2}{\sum e_b^2} e_b$<br>$= \frac{e_2}{d_2} - \frac{cov\left(\frac{d_2 e_1 + m_f e_2}{d_1 d_2 - m_2 m_f}\right)\left(\frac{e_2}{d_2}\right)}{\sum e_b^2} e_b$ |
| $M_{ca} = m_2 m_1$<br>$= \frac{m_2}{d_2} \frac{m_1 d_2}{(d_1 d_2 - m_2 m_f)}$ | $E_{ca} = m_2 e_1 + e_2 = \frac{m_2}{d_2} \frac{d_2 e_1 + m_f e_2}{d_1 d_2 - m_2 m_f} + \frac{e_2}{d_2}$ |

Prediction R1: When the feedback is negative 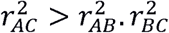

The reverse applies for positive feedbacks where 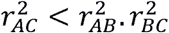

Prediction R2: In negative feedback *M*_*ca*_ > *M*_*cb*_*M*_*ba*_ and in positive feedback*|M*_*ca*_*|* < *M*_*cb*_*M*_*ba*_.

Prediction R3: The sign of this correlation will be decided by the sign of *m*_*f*_ which is negative for negative feedback and positive for positive feedback.

Prediction R4: (a) It can be shown that for negative feedbacks 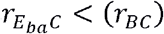 For positive feedback the prediction is more conditional. The inequality 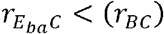 will be true above a threshold *r*_*BC*_. For smaller *r*_*BC*_ it is difficult to make a definite prediction. (b) for negative feedback 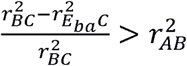 and for positive feedback 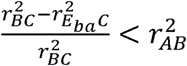 is true above a threshold *r*_*BC*_.

#### Positive or negative feed-forward pathway (P7)

The causal equations for this model would be

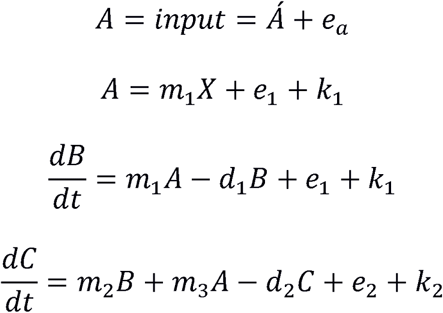

where *e*_*a*_*e*_1_*e*_2_ are not correlated. At equilibrium 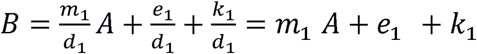 taking 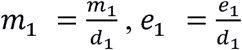 and, 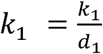

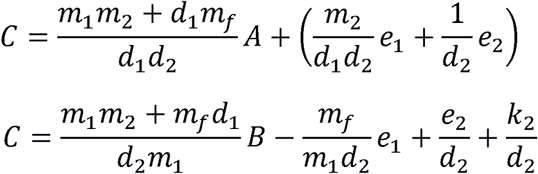

We will take 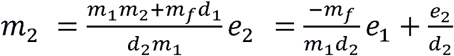

Note that since *e*_1_ decides both *e*_1_ and *e*_2_, the covariance between *e*_1_ and *e*_2_ will be 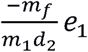, which will be positive when *m*_*f*_ is negative i.e. for negative feed-forward, and negative when *m*_*f*_ is positive i.e. positive feed-forward.

For simplifying the definition of feed-forward, we assume *m*_1_ and *m*_2_ to be positive, and the sign of *m*_*f*_ decides whether the feed-forward is positive or negative; a negative feed-forward pathway is once again associated with a problem of definition. If the feed-forward effect of *A* on *C* is stronger than that through *B,* and if their signs are opposite, the signs of slope in the causal and regression equations could be opposites. That is, if *m*_*f*_ *d*_1_ > *m*_1_ *m*_2_ then *M*_*cb*_ can be negative although the causal relationship between *B* and *C* is positive. This results in a paradoxical transformation of a causally negative feed-forward pathway into an effectively positive feed-forward pathway as the product *M*_*ba*_*M*_*cb*_ and *M*_*ca*_ both have the same sign.

Note that while all the expressions are the same as in feedback pathways (Table 9), the differences lie in the meanings of 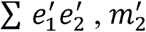 etc.

**Table 9.** Relationship between the causal and regression equations for Positive or negative feed-forward pathway.

| Slopes | Errors |
| --- | --- |
| $M_{ba} = m_1 = \frac{m_1}{d_1}$ | $E_{ba} = e_1 = \frac{e_1}{d_1}$ |
| $M_{cb} = m_2 + \frac{\sum e_1 e_2}{\sum e_b^2}$ $= \frac{m_1 m_2 + m_f d_1}{d_2 m_1} + \frac{\sum e_1 e_2}{\sum e_b^2}$ | $E_{cb} = e_2 - \frac{\sum e_1 e_2}{\sum e_b^2} e_b$ $= \frac{-m_f}{m_1 d_2} e_1 + \frac{e_2}{d_2} - \frac{\sum e_1 e_2}{\sum e_b^2} e_b$ |
| $M_{ca} = m_2 m_1 = \frac{m_1 m_2 + m_f d_1}{d_2 m_1} \cdot \frac{m_1}{d_1}$ | $E_{ca} = m_2 e_1 + e_2$ $= \frac{m_1 m_2 + m_f d_1}{d_2 m_1} \cdot \frac{e_1}{d_1} + \frac{-m_f}{m_1 d_2} e_1$ $+ \frac{e_2}{d_2}$ |

Prediction R1: For positive feed-forward 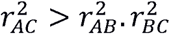, and for negative feed-forward 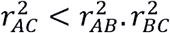, but under conditions in which the result mimics positive feedback, 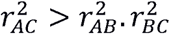 This is the condition when a causally negative feed-forward transforms into an apparent positive feed-forward.

Prediction R2: When *m*_*f*_ is positive, *M*_*cb*_*M*_*ba*_ *< M*_*ca*_ In the case of negative feed-forward *M*_*cb*_*M*_*ba*_ > *M*_*ca*_ but under conditions in which a negative feed-forward transforms into a positive feed-forward, *M*_*cb*_*M*_*ba*_ < *m*_1_ *m*_2_ = *M*_*ca*_.

Prediction R3: For positive feed-forward, we expect a negative correlation, and for negative feed-forward, a positive correlation. Therefore, for positive feed-forward, the correlation 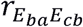 will have the opposite sign of that of *M*_*cb*_. For a negative feed-forward pathway, under conditions when it transforms into an effective positive feed-forward correlation, 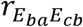 will have the opposite sign of that of *M*_*cb*_.

Prediction R4: (a) In the case of positive feed-forward 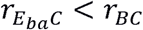 and in the case of negative feed-forward prediction is conditional. Under the conditions when a causally negative feed-forward becomes apparently positive feed-forward, the prediction of positive feed-forward is true. When a negative feed-forward is effective 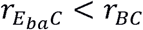 will be true above a threshold *r*_*AB*_. (b) For positive feed-forward 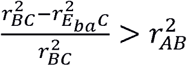. For negative feed-forward a universal prediction cannot be made. When the effect is that of a positive feedback the prediction of positive feedback is true, otherwise 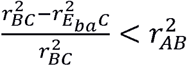.

#### Testing the null hypotheses

Testing in real life data needs to be different for equality and inequality predictions. The prediction can serve as the null model wherever equality is predicted, but needs to be treated as an alternative hypothesis wherever inequality is predicted. For pathways that predict equality, a two-tailed probability is used, and for pathways predicting one-way inequality, a one tailed test is used. In the results of simulations reported below, the convention consistently followed through Figs 2 and 3 is that if H_0_ is true, it is indicated by green, while H_1_ being true is indicated by red and H_2_ being true by yellow.

**Fig 2.**
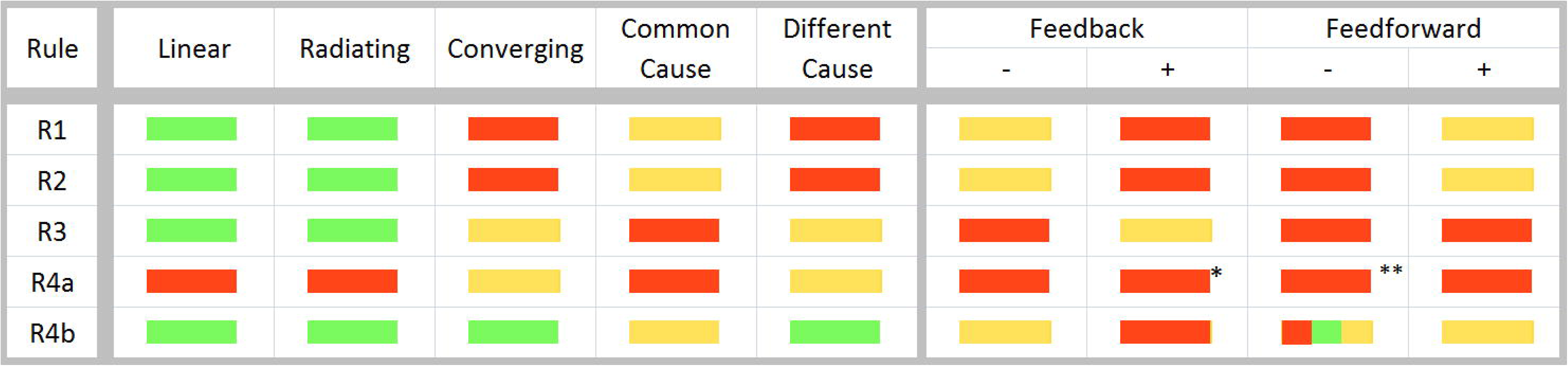
Analytical signatures for each pathway. Summarizing the analytical signature for each pathway in a color code where green represents acceptance of the null hypotheses H_0_, and red and yellow represent the acceptance of H_1_ and H_2_ respectively. Asterisks indicate conditional prediction e.g. 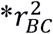 above a threshold, 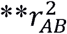 above a threshold.

**Fig 3.**
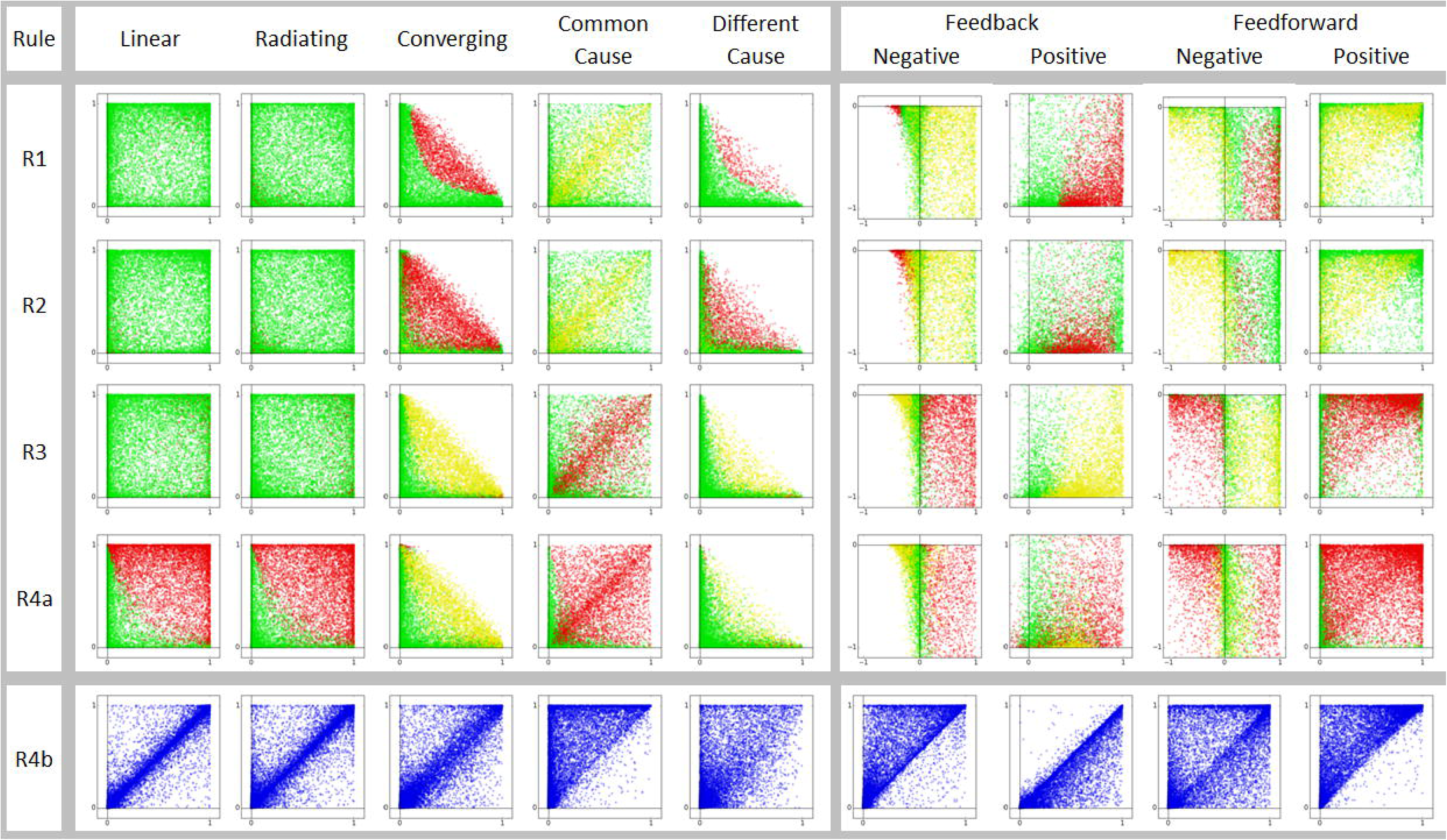
Simulation results of all pathways against all predictions. For all acyclic pathways and predictions from R1 to R4a, the result of every simulation run is plotted as a point with 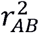 and 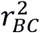 on X and Y axes respectively. Green represents null hypothesis true, red for H_1_ and yellow for H_2_. The results match very well with the predictions in Fig 2. For converging and different cause pathways, a pathway specific prediction is that the sum of the two correlation coefficients never exceeds unity. This is also evident in the simulation results.

R1: H_0_: 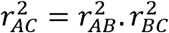 H_1_: 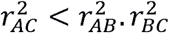 and H_2_: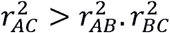. Since every correlation coefficient is associated with a confidence interval, to test the null hypothesis, we check that the confidence interval of 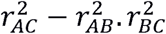 includes zero.

R2: H_0_: *M*_*ca*_ = *M*_*cb*_*M*_*ba*_ (green), H_1_: *M*_*ca*_ < *M*_*cb*_*M*_*b*_ (red) and H_2_: *M*_*ca*_ > *M*_*cb*_*M*_*ba*_ (yellow).

R3: Since the signs of causal slopes *m*_1_ and *m*_2_ are allowed to be positive or negative in the models, the correlation coefficients are multiplied by the sign of the slope *M*_*cb*_.

H_0_: 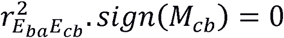 (green), H_1_: 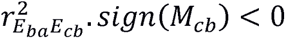 (red) and H_2_: 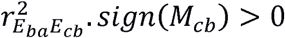 (yellow).

R4a: H_0_: 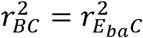 (green), H_1_: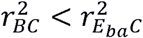 (red) and H_2_: 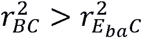 (yellow).

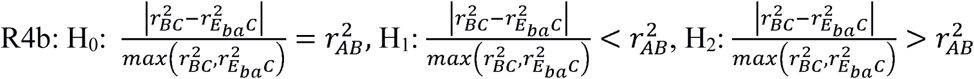. Since the prediction is about whether 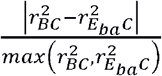 is predicted by 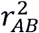, in the simulations results reported below we show a scatter plot between the two where good predictions lie along the diagonal and failure of prediction strays away from it.

For feedback and feed-forward pathways predictions from R1 to R4a, the X axis is *r*_*BC*_ and Y is 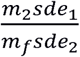 which reflects the relative strength of the feedback or feed-forward term as compared to the forward relation between B and C. The feed effect is strong when the ratio is close to zero and weak moving away from it. For negative feedback and feed-forward the Y axis goes from -1 to 0 and for positive feedback and feed-forward from 0 to 1. With negative feedback and feed-forward, there is an apparent conversion to positive feedback and feed-forward respectively under a set of conditions. When this happens *r*_*BC*_ becomes negative and the predictions of positive feedback and feed-forward respectively apply. It can also be seen that when the ratio is close to zero, predictions of converging pathway hold true.

Predictions from R4b for all pathways are shown as scatter plots with 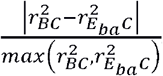 and 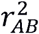. When they are predicted equal, most points lie along the diagonal. Wherever inequality is predicted, they are on one side of the diagonal.

Rejection due to overfitting inequality: For all inequality predictions, overfitting is possible. For example, if we expect that 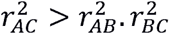, it is also possible that 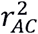 is too large than what can be predicted by the pathway under consideration. It is possible to test this either analytically using equations derived for the corresponding predictions (see S1 Text), or using simulations only if the parameters of the causal equations are known. If parameter estimates for the pathways are known from independent empirical sources, it should be possible to test over-fitting inequality. We will illustrate this with real life data later in the section ‘Testing specific pathways and questions: The case of pre-diabetes.’

### Simulations to test the sensitivity and robustness of predictions

We used simulations to test the sensitivity and reliability of the analytical predictions. The simulations were run using the causal pathway equations for each of the pathways *P1* to *P7* to generate data, assuming the errors to be distributed normally around a mean zero. Up to 10000 simulations are run, with each run using randomly drawn parameters and error standard deviations from a given range (see S1 Text ‘Simulations used for testing the sensitivity and robustness of the predictions’ for details). The error standard deviation ranges were selected such that the coefficients of determination were well spread between zero and one. The generated data were then used to test the predictions of the corresponding pathway. Simulations used in this section are not based on real life data, and are mainly employed to test the reliability of the predictions over a range of regression correlation coefficients.

### Agreement between analytical predictions and simulation results

Figs 2 and 3 show that simulations generally follow the analytical predictions quite well, but with certain limitations. Many of the predictions, particularly when H_1_ and H_2_ are expected to be true, work well above threshold values of *r*. When either 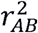 or 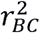 or both are small, the null hypothesis fails to get rejected. This threshold of sensitivity can be reduced by increasing sample size (n) (Fig 4). In the case of cyclic pathways, many predictions are conditional as described above, and that is clearly reflected in the simulations. The agreement between predictions and simulation results is weaker for a few specific pathway-prediction combinations, in the sense that they work in a narrow range of conditions. This was seen in case of P5 (different cause) prediction from R4b, and P7 (negative feed-forward) prediction from R2. The predictions become more reliable at higher n. This implies that we need to be conservative in rejecting pathways in such pathway-prediction combinations, particularly when the correlation coefficients are small. Further, wherever the predictions are themselves the null models, its rejection will naturally become conservative at low correlations. However for inequality predictions, where the null hypothesis is equality, failure of rejecting the null hypothesis should not be taken as rejection of the prediction when correlations are weak. When we take such a conservative approach, rejection of a prediction can be confidently taken to mean rejection of a pathway.

**Fig 4:**
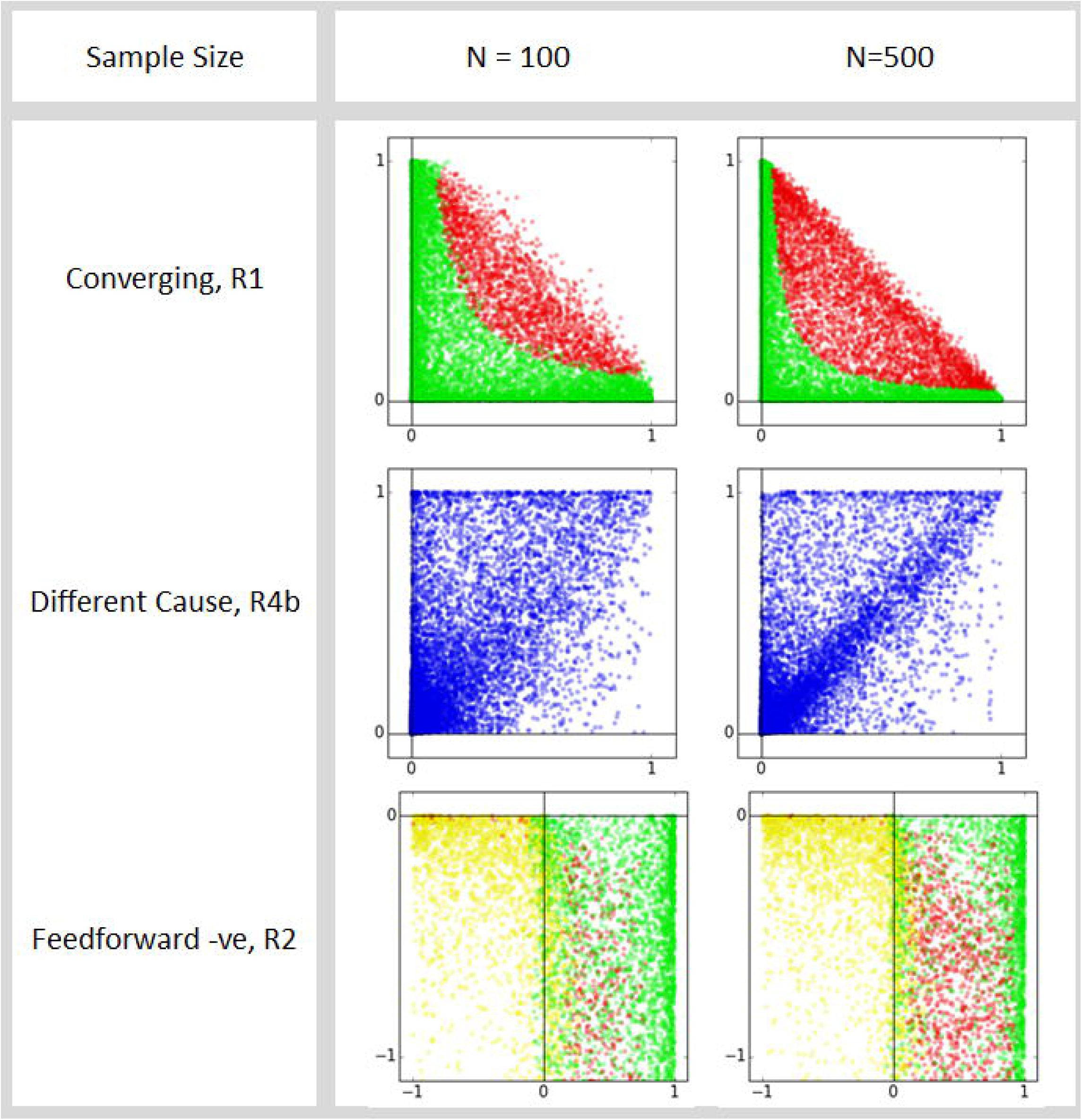
Effect of n on the reliability of prediction. Note that the parameter area over which the simulation results match the prediction increases with n.

It can be seen from Table 10 that each pathway makes a set of predictions by which some pathways can be differentiated from others. However, some have an identical set of predictions among the general predictions described so far. Table 10 shows that there are 6 different signatures among 9 primary pathways. Some of the predictions are conditional, and therefore, it may not always be possible to differentiate between pathways. For example, some predictions do not work for very small *r*^*2*^ values, or if feedback is not distinguishable from linear pathways unless the feedback arrow is sufficiently strong. Such limitations are common to all statistical tools, and they need to be used and interpreted in light of the appropriate context and conditions.

**Table 10.**
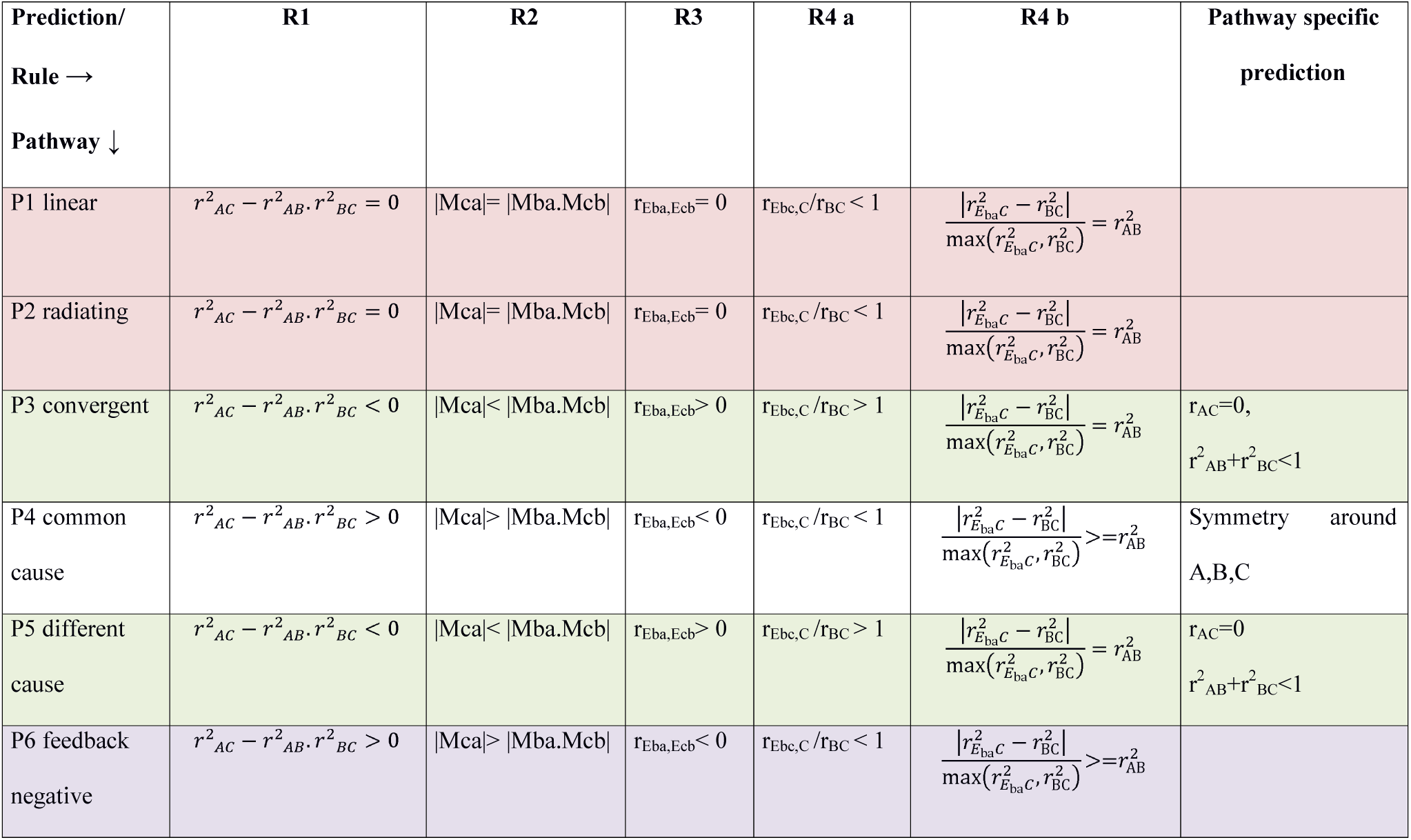
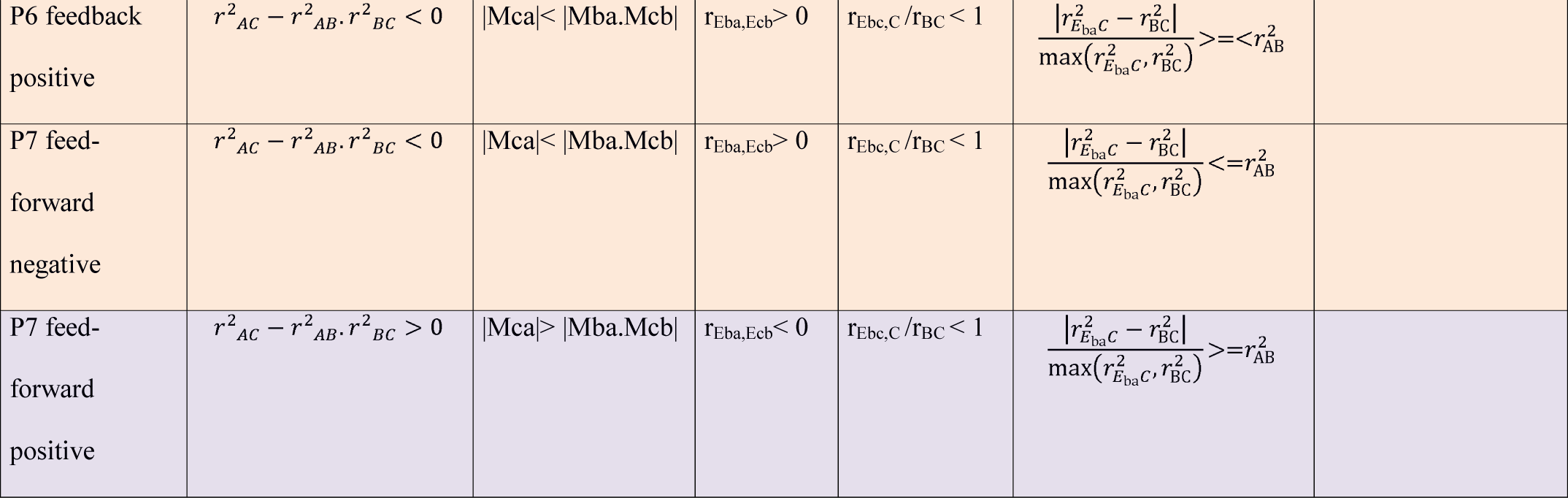
Summary of predictions of all pathways considered.

Table 10 legend: Note that there are 6 distinct signatures among 9 pathways. Pathways with identical predictions are shaded with the same colour. There is some redundancy between the predictions. For example the results of R1, R2 and R3 are tightly correlated. We feel that the redundancy serves to validate and reinforce the predictions. Also when there are complex pathways arising through combinations of primary pathways, different predictions show differential departures from the primary pathway predictions. Therefore all predictions are useful in spite of some redundancy.

### Sensitivity analysis

The predictions derived and tabulated above are based on the typical assumptions of mainstream statistics that the input variables are distributed normally and that all causal links are linear. However, it is important to ask how critical these assumptions are for the predictions to work. In experimental biology, the input variable is often designed to have uniform intervals and is not normally distributed. A moderate deviation from linearity is also common in physiological and other biological systems. If the predictions are too sensitive to these assumptions, they may prove to be of limited use in real-life. We used Monte-Carlo simulations to assess whether the predictions work under moderate deviations from these assumptions. When the input variables were selected randomly from a uniform rather than a Gaussian distribution, all predictions worked with nearly the same differentiating ability (data not shown). Similarly, when non-parametric Spearman ranked correlations were used instead of Pearson’s correlations, the correlation related predictions (from R1 and R4) worked similarly (data not shown). This demonstrates that the tools are not too sensitive to the assumptions of normality of input variable, linearity of relationships, and parametric or non-parametric nature of correlations.

### Applications of the method

#### Accepting or rejecting pathways using real-life data

Two approaches are possible by which the predictions of a pathway can be tested using real-life data. (i) Based on confidence intervals of correlations and regression slopes: The null hypotheses for every prediction can be tested using calculation of confidence intervals of regression correlation parameters. Simulations have shown that except when the underlying correlation coefficients are too low, this approach can be reliably used to test the predictions. The sensitivity of predictions depends upon the sample size as well as the position in the parameter space (Fig 3). It is likely therefore that at smaller sample sizes, or at lower 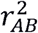 or 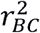, pathways that predict H_1_ or H_2_ may fail to get support even if true. On the other hand, at lower 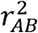 or 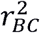 if a pathway predicts H_0_ to be true and the null hypothesis gets rejected, the rejection can be highly reliable. (ii) Monte-Carlo simulation approach: An alternative approach, which will be more conservative in rejecting pathways, is the Monte-Carlo approach. Assuming a specific pathway to be true, it is possible to back calculate the causal equation parameters from the regression correlation parameters obtained in the data (Tables 2 to 9). For pathways such as negative or positive feedback, it is not possible to estimate all causal parameters from regression parameters. In such cases, if empirical estimates of one or a few causal parameters can be obtained, the remaining causal parameters can be worked back. Using the estimated parameters of causal equations, Monte-Carlo simulations can be run to obtain the probabilities of getting the observed results. This approach can be particularly useful when the correlations obtained in the data are weak, and a conservative inference is preferred.

#### Distinguishing between pathways with identical signatures

From the predictions summarized in Table 10, it can be seen that some pathways share prediction signatures. For example, the linear pathway cannot be distinguished from radiating or convergent, is indistinguishable from different cause. There are three possible ways of resolving between pathways with similar signatures: (i) Swapping variables: In the generalized predictions, common cause pathway and negative feedback pathway have the same predictions. However, the predictions of the common cause pathway are symmetric around A, B and C, and flipping the positions of the three variables does not alter the predictions, which is not the case with negative feedback pathway. (ii) Involving a fourth variable whose causal relationship with at least one of the triad is already known, or (iii) involving more variables to cross validate pathways. We will discuss (ii) and (iii) in a different context below.

#### Inferring causality between two variables

It is extremely difficult to infer causal relationship between two correlated variables. Although some solutions have been suggested, their applicability is limited (15, 16). However, it is possible to infer the causal relationship between two variables if we have data on a third variable that is correlated with one or both of them with known causality. For example, in men, testosterone levels and muscle strength are correlated, but the direction of the causal arrow might be uncertain since testosterone can increase muscle mass, while (34) exercise can also induce a testosterone response (35,36). The causal relationship can be revealed in cross-sectional data if we use chronological age as a third variable. Neither testosterone nor muscle mass decides the chronological age, but age may affect one or both the variables. If age shows significant correlation with one or both the variables, the predictions from different possible pathways can be tested using the set of predictions as described. By testing these predictions, it should be possible to determine the causal relationship between muscle mass and testosterone.

#### Inferring causal pathways with three variables

To infer causal pathways within three intercorrelated variables, three alternative approaches are possible. The first approach is to test and resolve between preconceived hypothetical pathways. It is likely that prior knowledge or some insights into mechanisms allow us to start with a few plausible alternative pathways. It is possible to perceive more complex pathways by combinations of the primary pathways that we considered in this paper. For example, a pathway may contain both feedback and feed-forward elements. Such complex or combinational pathways can be used to make a set of predictions by the analytical approach described above, and testing these predictions can resolve between pathways. If we do not have such preconceived pathways, it would be necessary to consider all possible combinations of pathways between the three variables, and make differential predictions from each of them. In such cases, we must also consider permutations of the variables. At the end, it may not be possible to ascertain a single unique causal pathway since the prediction signatures of some of them may be identical. Nevertheless, it would still be possible to reject some pathways based on their prediction signature. In addition, if available, we can involve a fourth variable correlated to one or more of the three, if there is some pre-existing knowledge about its causal relationship.

#### Inferring causal networks with more than three variables

In complex systems, often there are large causal networks. In such networks, combinations of 3 membered motifs can be identified. Out of the possible pathways among three variables some can be rejected using analysis of the three variables. Bringing in a fourth one can provide additional insights which can be used for cross checking or validating our first set of inferences. In complex causal networks, there can be many such cross check and validation possibilities. For large networks algorithms requiring massive computational power may be needed that may pin down one or a few network structures from the large number of possible ones using combinations of three member motifs and cross validation facility among the motifs.

### Testing specific pathways and questions: The case of pre-diabetes

Apart from some common pathways described above, it is possible that real life problems have some added complications due to which, the standard solution of testing a fixed set of predictions may not be sufficient. However, one can apply similar foundational principles to handle such pathway-specific questions. We will illustrate this using a classical hypothesis that attempts to explain a human physiological state designated as an insulin resistant, hyperinsulinemic, normoglycemic, pre-diabetic state. In this state, the plasma levels of fasting insulin (FI) are raised although fasting glucose (FG) remains normal. The classical interpretation of this state (Fig 5a) is that a rise in insulin resistance, presumably as a result of obesity, is primary. Insulin resistance interferes with insulin-induced glucose uptake by muscle and other insulin-dependent tissues. The reduced uptake raises plasma glucose levels. The raised plasma glucose induces extra insulin secretion so that plasma insulin levels go up. The extra insulin compensates for insulin resistance and normalizes glucose level. As a result, the fasting steady state of an insulin-resistant individual is characterized by raised FI and normal FG. At steady state, insulin resistance is measured by the index, HOMA-IR (defined as 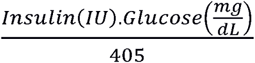), and the β cell response to glucose, by the index, HOMA β (defined as 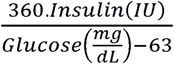). Both the indices are based on the assumption of a steady state.

**Fig 5:**
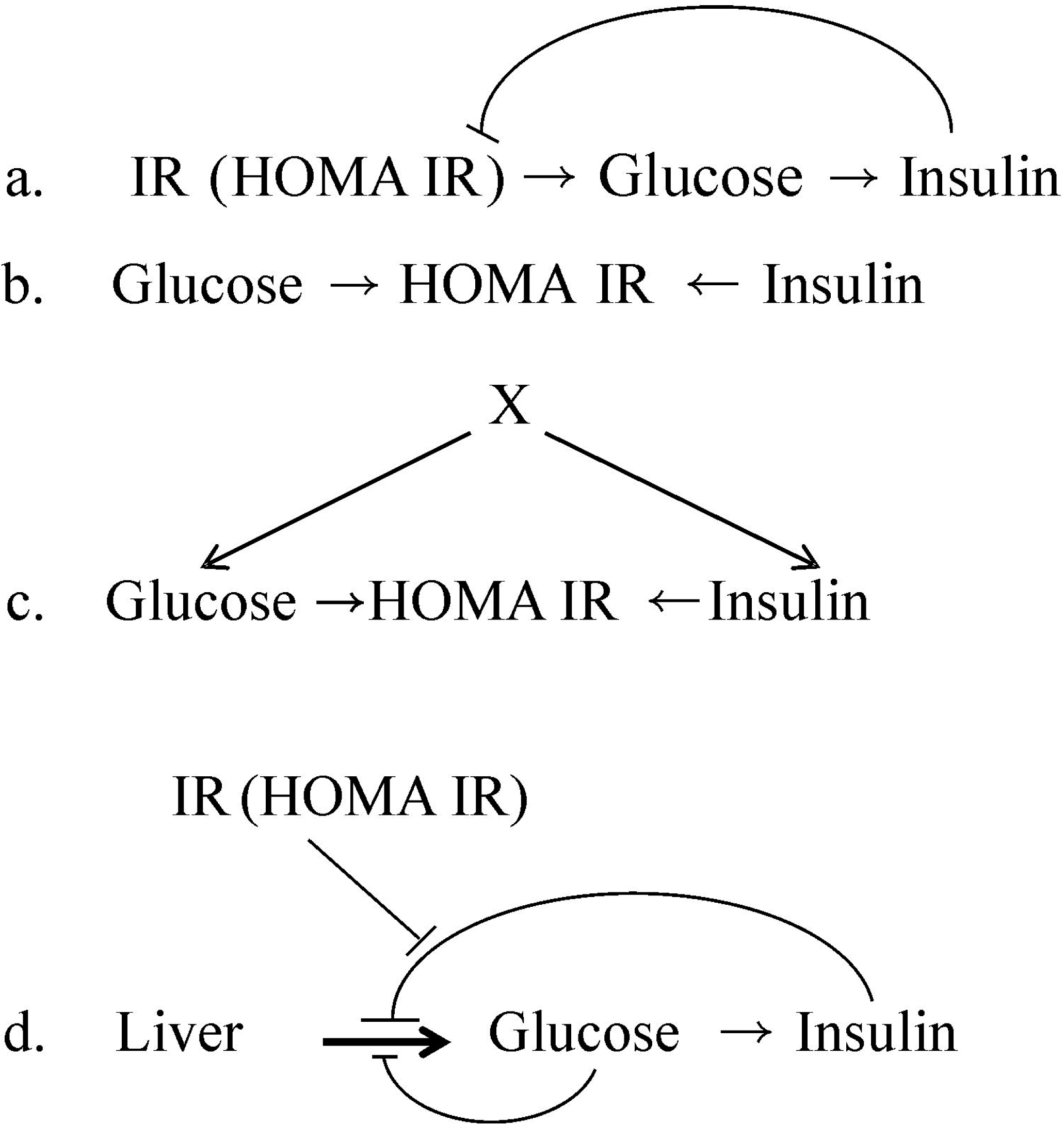
Possible pathways between insulin resistance, FG and FI: a) A simplified single feedback pathway that approximates the negative feedback pathway P6. b) A null model assuming FG and FI to be independent and HOMA-IR a derived construct. c) An improvised null model with an external causal factor influencing FG and FI. d) The classically perceived pathway with dual feedback from glucose and insulin.

A logical flaw in this interpretation is that, after the glucose levels return to normal, there is no reason why FI should remain high. Insulin has a short half-life of about 6 minutes (32,37) and therefore a steady state level can be achieved quite fast; 12 hour fasting should be sufficient to achieve such a steady state. Therefore, a steady state in which FI is raised but FG remains normal is not well explained by the classical theory. In spite of this flaw, the main stream thinking in this field has held on to this interpretation for over four decades, and the indices HOMA-IR and HOMA-β continue to be commonly used in epidemiological research. Challenges to this causal interpretation come from the arguments and evidence that rise in FI precedes insulin resistance (24–28,38). Therefore, there is a need to reexamine the classical causal pathway. We will test this pathway based on our interpretations of the interrelationships of the regression-correlation parameters.

The pathway in question is more complex than the basic set of pathways P1 to P7. For regulation of glucose production by the liver and glucose uptake by tissues, there is a dual negative feedback. One feedback is exerted by glucose itself, which enhances tissue uptake and suppresses liver glucose production. The other feedback operates through insulin, which facilitates glucose uptake by insulin-dependent tissues and suppresses liver glucose production. If we ignore the direct glucose feedback and assume that feedback regulation operates only through insulin, then there is a single negative feedback. Thus the pathway can be simplified to the negative feedback pathway P6. If we incorporate dual feedback, as the equations show below, the relationship between insulin resistance and FG is not strictly linear. We could therefore use the standard set of predictions of a negative feedback model, assuming a single feedback. Alternatively, we can use the dual feedback model, and apply simulations to make and test predictions, since empirical estimates for most of the parameters are available from experiments (see S2 Text).

However, the main problem in testing these pathways is that we have no direct measure of insulin resistance. HOMA-IR and HOMA-β are believed to measure insulin resistance and β cell response respectively, but they are derived from the other two variables, which makes the problem tricky and circular. We approach the problem using more than one set of assumptions. (i) First, we test the dual feedback pathway (Fig 5d) assuming HOMA-IR and HOMA-β to faithfully represent insulin resistance and β cell response respectively. (ii) Then, we examine the constraints laid down by deriving these two parameters from the other two variables. (iii) In comparison, we use a null model (Fig 5b) in which the classical pathway is not true, there is no relationship between FG and FI, and HOMA-IR and HOMA-β are artificial constructs derived from the two measured variables and may not reflect any real phenomenon. We also test the typical convergent model, in which FG and FI determine HOMA-IR. (iv) Using some oversimplification, ignoring non-linearity of the model and assuming that HOMA-IR and HOMA-β are faithful indicators, we test the classical predictions of the negative feedback pathway (Fig 5a) as described earlier. We use epidemiological data on FG and FI measurements in four populations to test the classical causal pathway using our approach.

#### Data sources

We used four data sets of sample studies by two research groups. All the four sets contain individuals with and without overt type 2 diabetes. Since we are addressing the prediabetic state here we have taken the non-diabetic subset of *n* individuals from the four samples. (i) Coronary Risk of Insulin Sensitivity in Indian Subjects (CRISIS) Study, Pune, India (39) (n=558). (ii and Newcastle Heart Project (NHP), England, (40) which has data on populations of two different ethnic origins namely European white (n=595) and south Asian (n=413). (iv) Pune Maternal Nutrition Study (PMNS), Pune, India (41) (n=299). All the predictions are tested independently in all the four data sets.

#### The dual feedback model (Fig 5d)

We assume that the standing plasma glucose level is a result of baseline rate of glucose production by the liver; suppression of this production as well as muscle glucose pickup which is proportional to the standing glucose level (direct glucose feedback); the insulin mediated suppression as well as uptake (insulin mediated feedback) and individual variability. The standing insulin levels are a result of glucose stimulated insulin secretion on the one hand and insulin degradation on the other. Thus, the causal equations can be written as

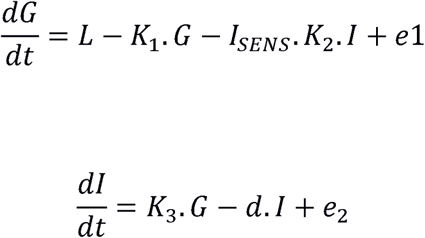

Where G and I are plasma levels of glucose and insulin respectively, and FG and FI are the fasting steady state levels of the same. *K*_*1*_ denotes the rate constant for negative feedback of glucose on liver glucose production and tissue glucose uptake; *K*_*2*_ denotes the rate constant for insulin-mediated feedback which is proportional to *I*_*SENS*_, the insulin sensitivity of tissues; *K*_*3*_ is the rate constant for glucose-induced insulin release; and *d,* the rate of insulin degradation.

Steady state solution: By assuming the net change to be zero in a steady state, we get

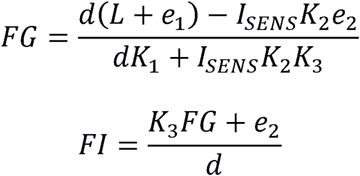

It can be seen that the steady state glucose level is a function of insulin resistance (*IR*) which is a reciprocal of insulin sensitivity. Using the reciprocal, we can write

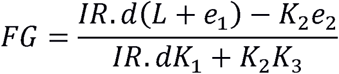

Thus the relationship between FG and IR is non-linear and follows a saturation curve. Testing the pathways by the four different approaches described above:

i. Assuming classical pathway and faithful indices: The following predictions of the classical pathway depicted in Fig 5d and modeled above are testable.
  a. HOMA-IR, FG and FI should be positively correlated to each other. This prediction is true in all the four data sets except that the correlations between FG and FI are weak in all the four data sets. In terms of the variance explained (range 2.6 to 4.9 %) FG and FI are poorly related (Table 11a). The glucose homeostasis model expects a positive correlation between FG and FI. It is important to realize this since in the classical thinking, a prediabetic state is characterized by increased insulin but normal glucose levels. If the compensatory insulin response is mediated through glucose, it is impossible to have a raised FI without a proportionate rise in FG. In the pathway predictions, a positive correlation between FG and FI is expected independent of the feedback loop. However the classical thinking tries to explain a hyperinsulinemic normoglycemic state achieved through this pathway. The poor correlation between FG and FI, and a large coefficient of variation in FI compared to FG indicates that a normoglycemic hyperinsulinemic state may indeed be achieved, but whether the classical pathway offers a sound explanation for this state is the question. In an insulin resistant state, the level of FI can increase by about 10-fold the normal. However, the difference between the lower and upper limit of glucose in a pre-diabetic state is less than 1.5-fold. To achieve a tenfold increase in the effect resulting from a 1.5 fold increase in the causal variable, the slope needs to be of the order of 7 to 8. However in the data, the regression slope ranges between 0.05 and 0.2 (Table 11). Therefore the variance in FI is unlikely to be caused by variance in glucose following insulin resistance. Therefore, we need to conclude that most of the variation in FI appears to be random error independent of insulin resistance.

**Table 11.**
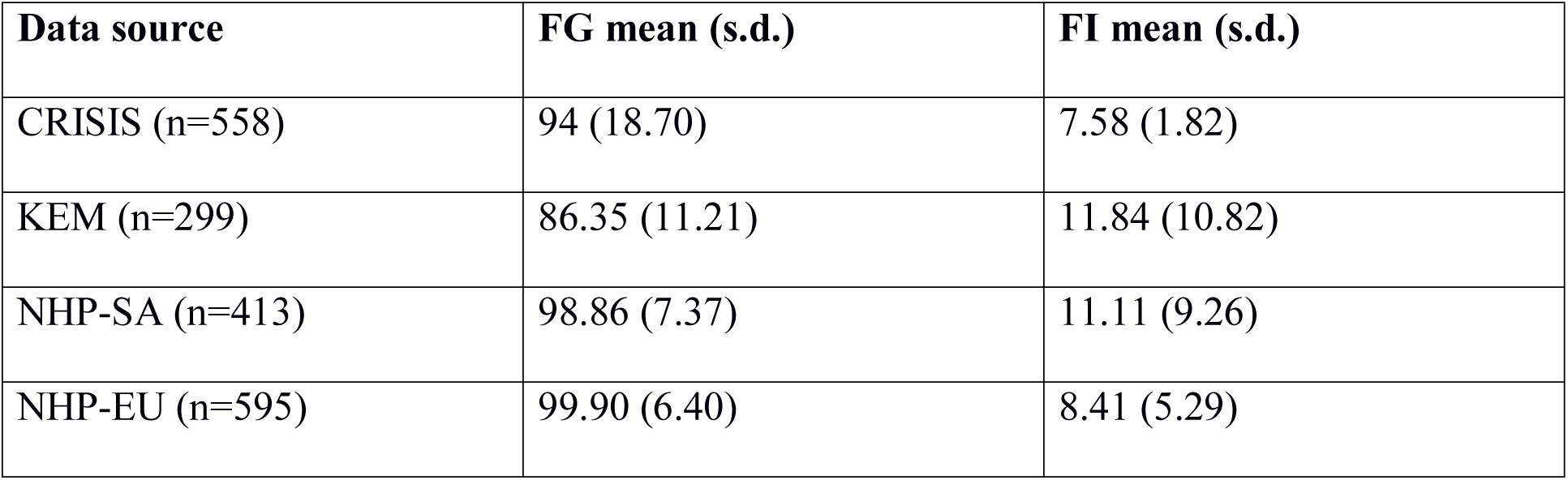
Testing the putative pathway leading to a hyperinsulinemic, normoglycemic, insulin resistant prediabetic state. **a. Fasting glucose and fasting insulin in the four datasets**

| Data source | FG mean (s.d.) | FI mean (s.d.) |
| --- | --- | --- |
| CRISIS (n=558) | 94 (18.70) | 7.58 (1.82) |
| KEM (n=299) | 86.35 (11.21) | 11.84 (10.82) |
| NHP-SA (n=413) | 98.86 (7.37) | 11.11 (9.26) |
| NHP-EU (n=595) | 99.90 (6.40) | 8.41 (5.29) |

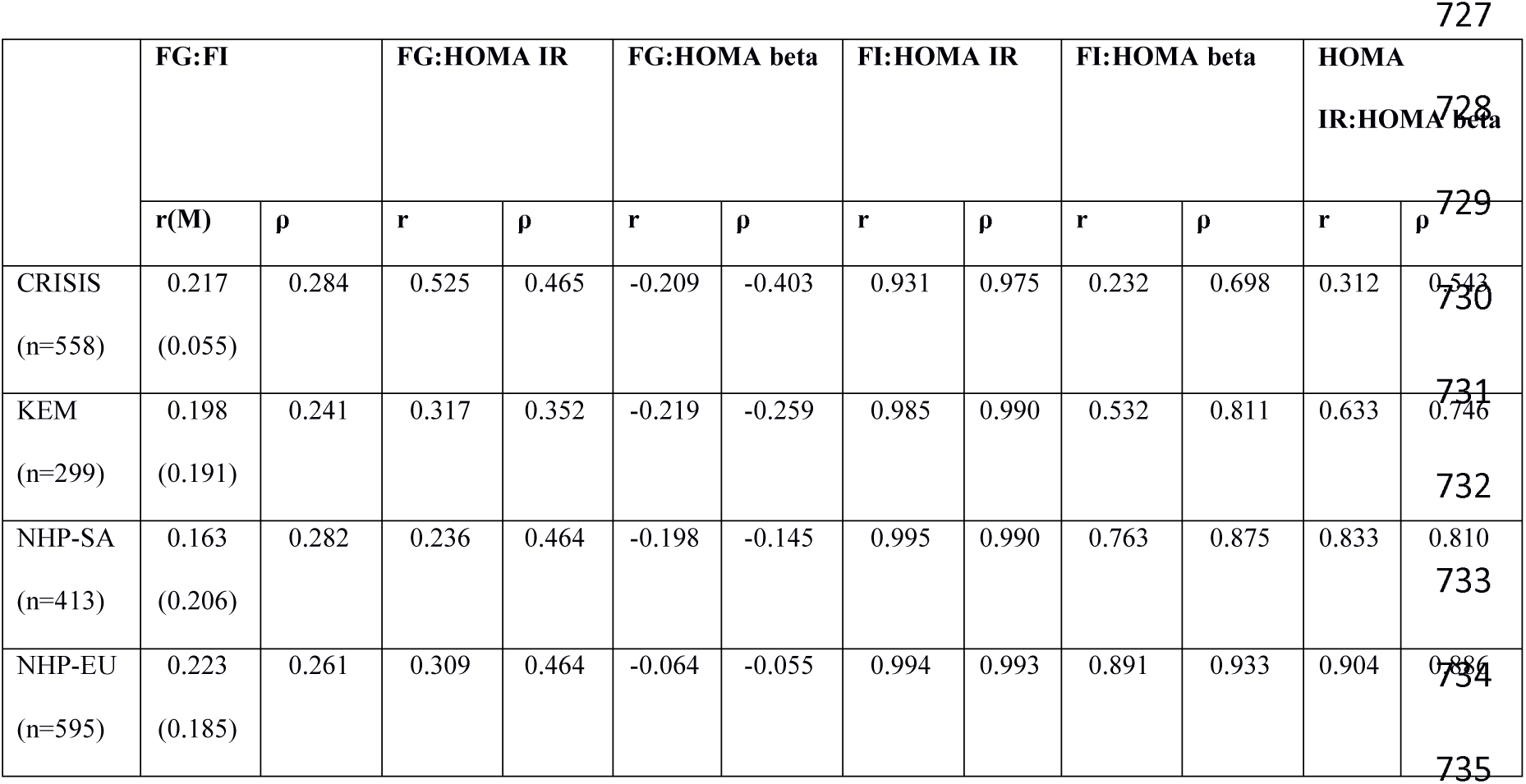
**b. Correlations between FG, FI, HOMA-IR and HOMA beta in the four data sets**

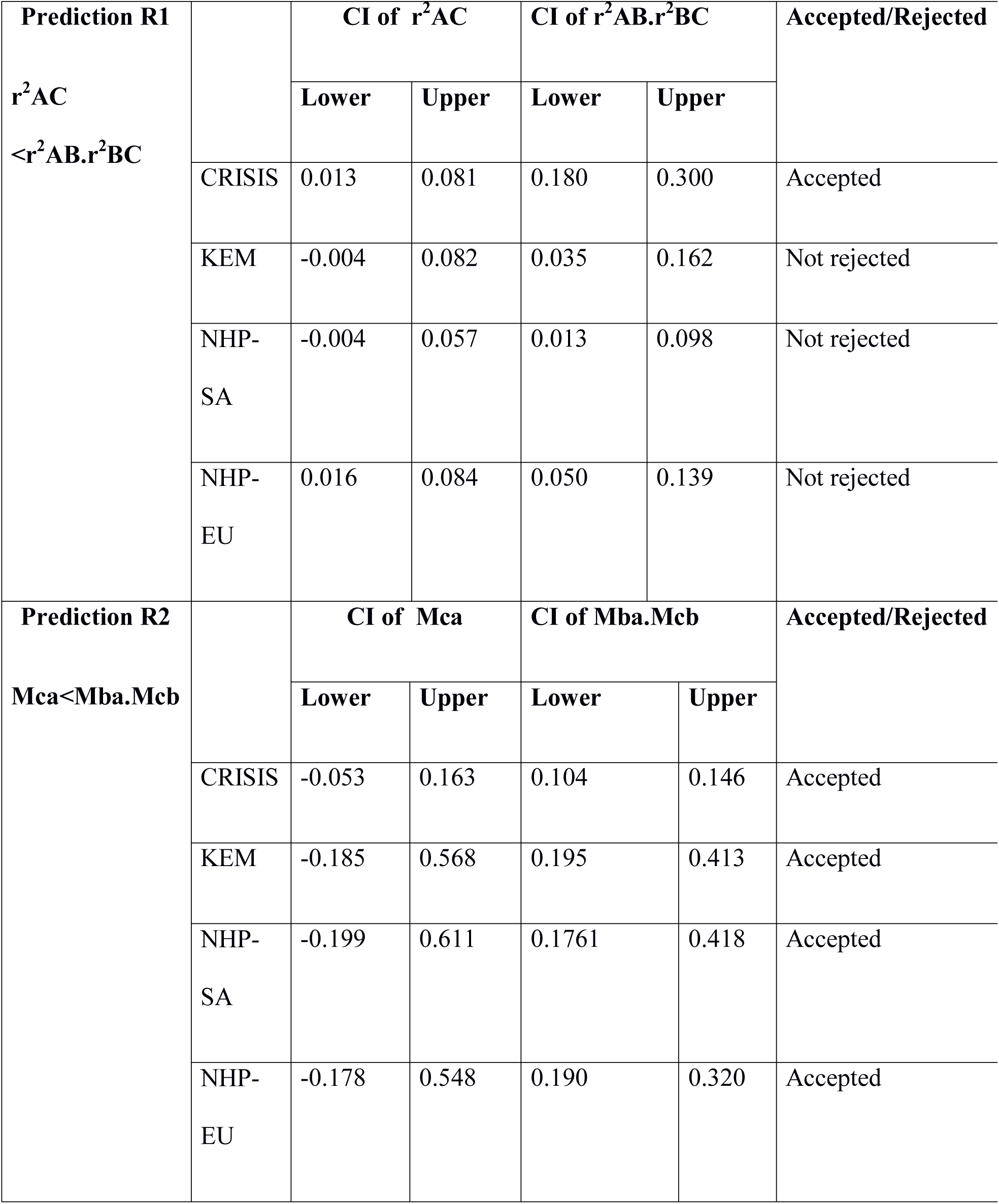
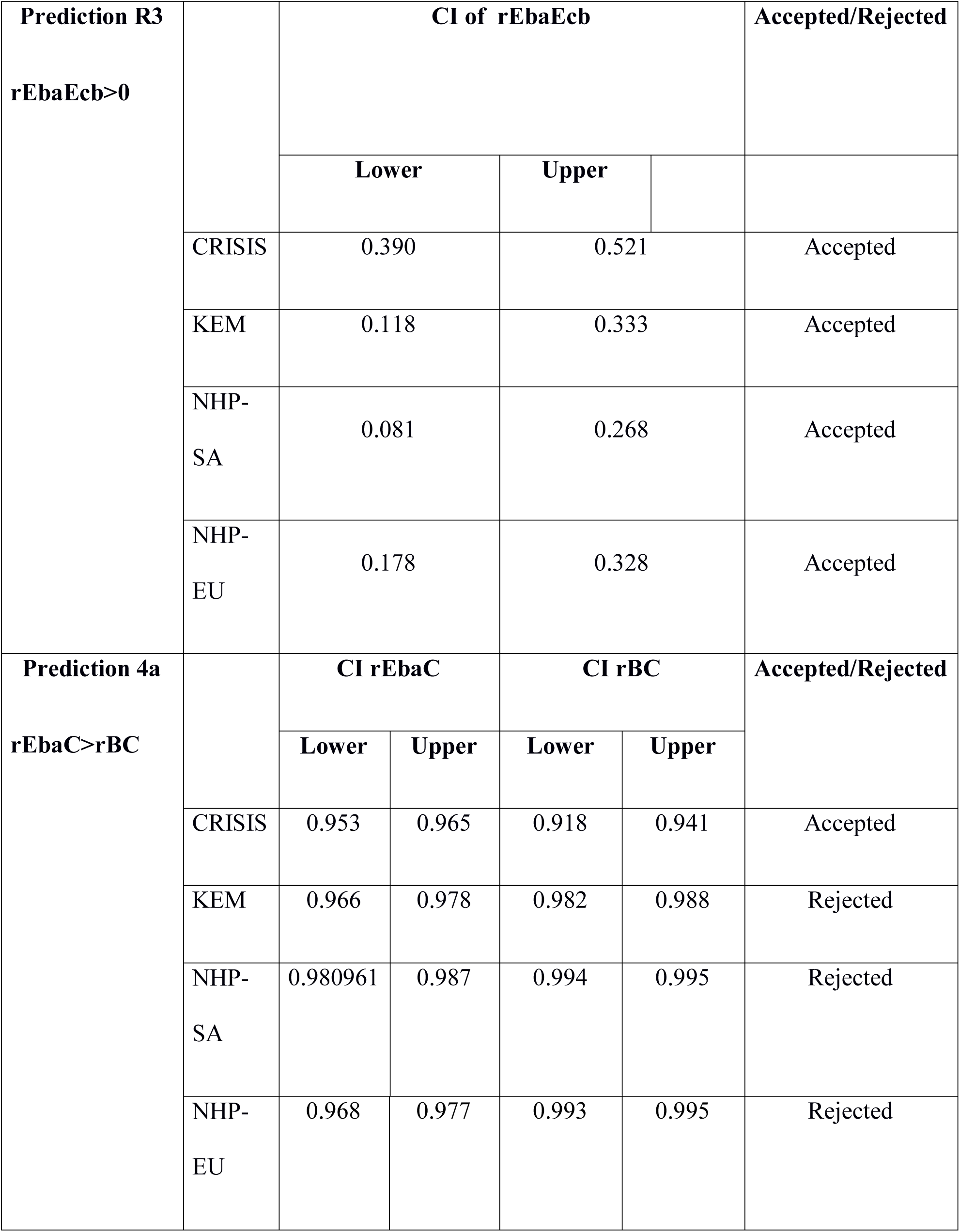
**c. Convergent pathway (A=Glucose, B=HOMA IR, C=Insulin)**

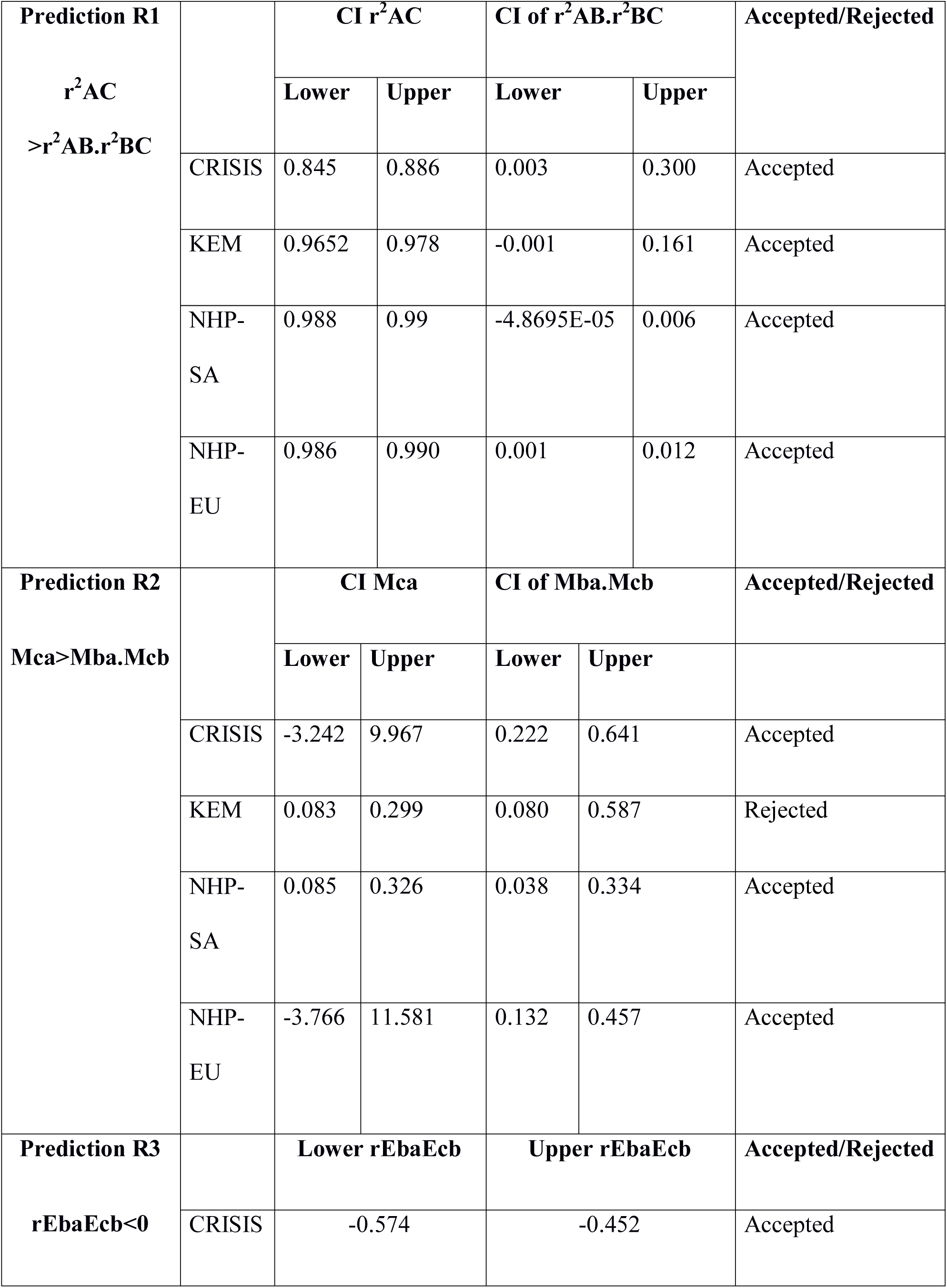
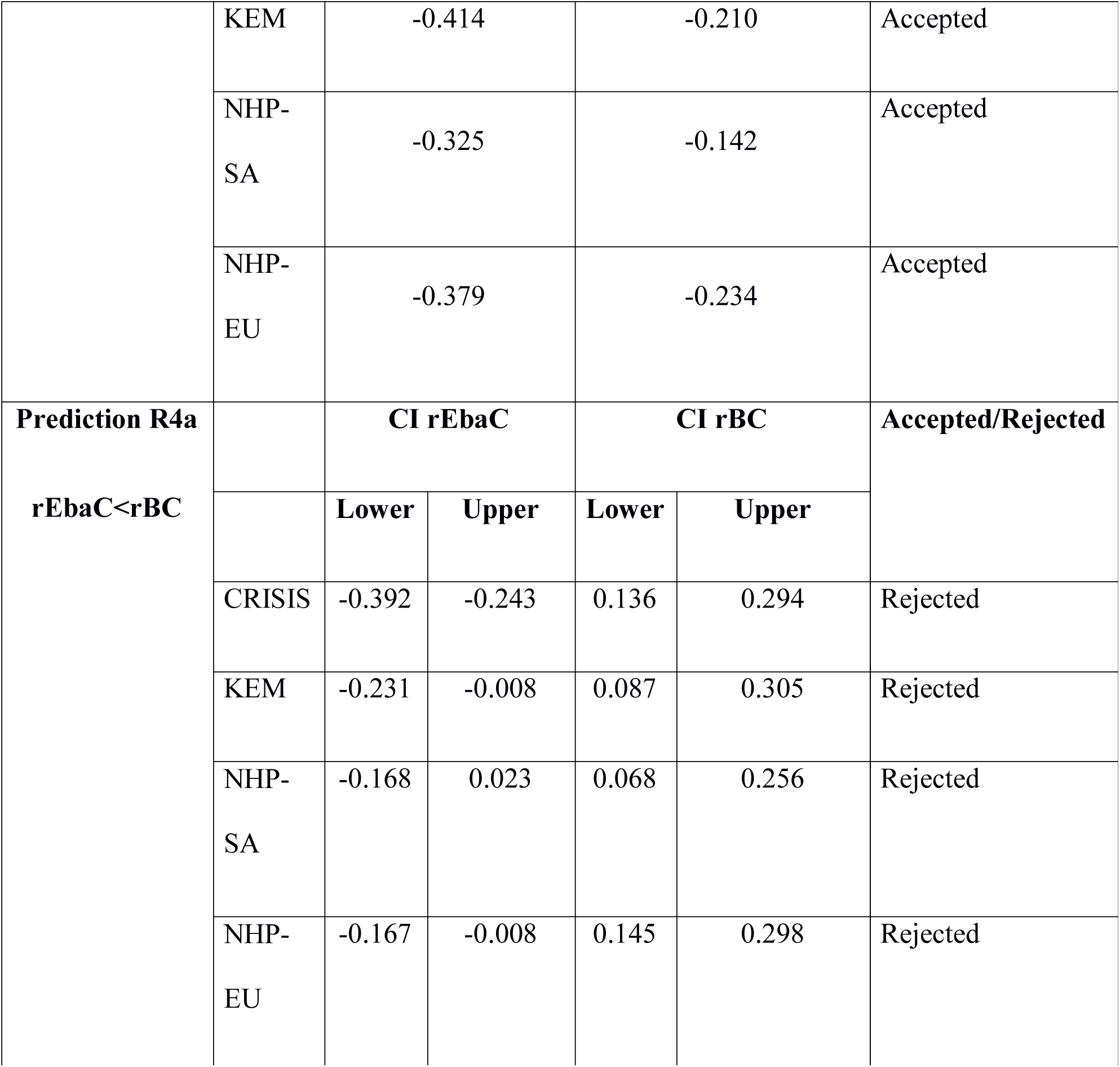
**d. Negative feedback pathway (A= HOMA IR, B=Glucose, C=Insulin)** Table 11 footnote: The classical pathway leading to a prediabetic state is tested using the pathway prediction approach. (a) Means and standard deviations from the four data sets (b) correlations obtained from empirical data (c) testing the four predictions for the null model (fig 5b) (d) testing the four predictions of a simplified classical negative feedback pathway (fig 5a).
  b. By the steady state equations, the slope of the regression of FI on FG should be *K*_*3*_*/d*. Empirical estimates for both *K*_*3*_ and *d* are available (see S2 Text) and therefore this prediction can be tested. The empirical estimates are *K*_*3*_ = 0.08 microIU.mg/min and *d* = 0.15/min respectively, and thereby the expected slope is 0.533. In all the four data sets, the slopes are significantly smaller than the ones predicted from the empirical estimates (0.05 to 0.2). Thus, apart from a mismatch between the slope required to cause the observed variation in FI and actual slopes, the slopes expected from the empirical estimates of parameters and those obtained in regression also do not match. The latter mismatch by itself may not be sufficient to reject the pathway since a large measurement error in the X variable, i.e. FG can lead to underestimation of regression slope, but this explanation implies that a substantial part of variation in glucose is independent of insulin resistance, and is akin to random error with respect to the hypothetical causal pathway.
  c. HOMA-β in our assumption represents *K*_*3*_. However *K*_*3*_ is a constant in our model, and although it may have some variability in the population, it is uncorrelated with the three variables of concern. Therefore, HOMA-β should show no significant correlation with FG, FI and HOMA-IR. However, in all the four data sets HOMA-β is significantly positively correlated with FI, but negatively correlated with FG and positively correlated with HOMA-IR.
  d. In a negative feedback pathway 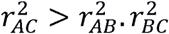. Qualitatively this inequality is true for HOMA-IR, FG and FI in the data. However, simulations show that there is overfitting of the inequality. 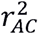 in all the four sets of data are substantially higher than the distribution obtained in the simulations (Fig 6). The correlation between FI and HOMA-IR is far greater than that predicted by the simulations, leading to an overfitting rejection.

**Fig 6:**
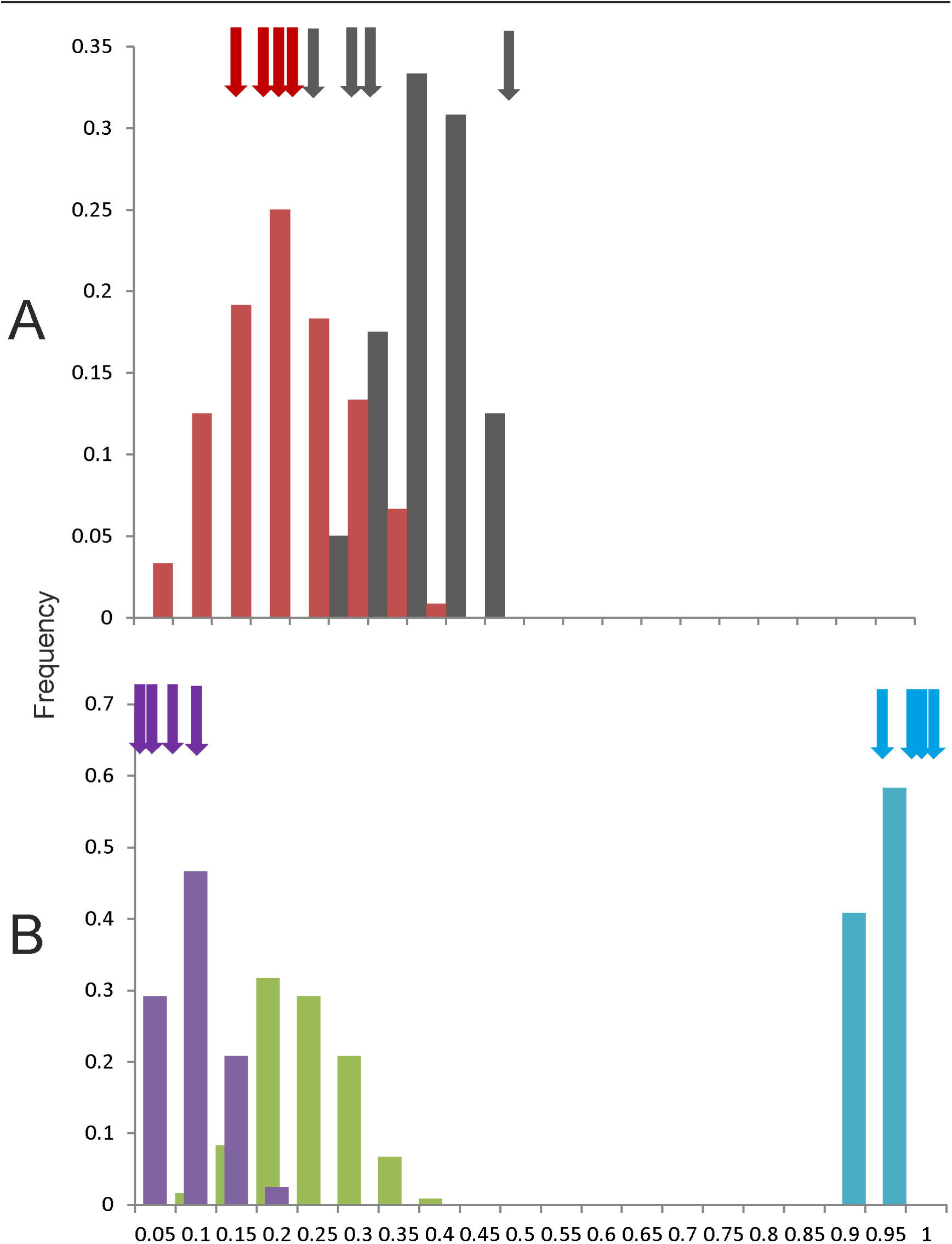
Frequency distribution of correlation coefficients in simulations of the classical pathway leading to prediabetic state: Bars represent the distribution of Pearson’s correlations obtained in 10000 runs of simulations. The arrows indicate Pearson’s correlations in the four sets of empirical data. The distribution generated by simulations matches well with the real life correlations for true IR-FG (grey bars and arrows), FG-FI (red bars and arrows), and the product of the two (purple bars and arrows). The correlation between true IR and FI is greater than the product as predicted by the pathway (green bars, we do not have empirical estimates of these correlations) but the correlation between HOMA-IR and FI (blue bars and arrows) is substantially greater than the predicted leading to an overfitting rejection. This indicates that either HOMA-IR as currently calculated is substantially different from true insulin resistance or the pathway get rejected based on this prediction. Thus if we assume the two HOMA indices to faithfully represent insulin resistance and beta cell response respectively, then classical pathway needs to be rejected owing to mismatches with many of its predictions.
  e. Effects of deriving HOMA-IR and HOMA-β from FG and FI: Since HOMA-IR and HOMA-β are not independently measured but derived from FG and FI measurements, some correlations will follow from the derivations themselves. The overfitting anomaly observed above can be explained as an artifact coming out of the calculation of HOMA-IR. However, some other anomalies remain unexplained. Here we are assuming that the classical pathway is true and therefore, FI is a linear function of FG. If FI is represented as *m.FG + e*, HOMA-IR will be correlated to FG^2^. Similarly, HOMA-β should be represented as *m.FG/(FG – 63)+e.* Under normal physiological range, FG > 63 and therefore HOMA-β is a decreasing function of FG. As a result both FI and HOMA-IR should be negatively correlated to HOMA-β. Simulations of the pathway results in a negative correlation between HOMA-IR and HOMA-β as long as the errors are small to moderate. These expectations do not match the empirical data, in which FI and HOMA-IR have significant positive correlations with HOMA-β. Thus, accepting the classical pathway with some allowance for artifacts coming out of the derived variables is not sufficient to explain the empirical correlations.
  f. Testing the predictions of the null model: If FG and FI are independent of each other and have some variance around a mean, HOMA-IR is expected to be positively correlated with both since it is a product of the two. FI should be positively correlated with HOMA-β, but FG should be negatively correlated with HOMA-β. In the HOMA-IR-HOMA-β relationship, FI is in the numerator of both. FG is in the numerator of HOMA-IR but in the denominator of HOMA-β. Nevertheless, since the coefficient of variation of FI is substantially greater than that of FG, FI is expected to dominate the relationship and result in a positive correlation between HOMA-IR and HOMA- β. All these predictions are observed in the data. The mismatch of the null model with the data is that it assumes FG and FI to be independent and uncorrelated. In all the four sets of data, there is a significant but weak correlation between the two. The r^2^ ranges from 0.026 to 0.049, and thus not more than 5 % of variance is explained by the relationship. If we consider FG and FI to be independent and HOMA-IR and HOMA-β derived from them, they constitute a convergent pathway that can be tested by the pathway predictions. It can be seen that predictions from R1, R2 and R3 of the convergent pathway are accepted. However, prediction from R4 and the pathway-specific prediction are rejected (Table 11b). These rejections can be explained by the positive correlation between FG and FI. We have seen in the analysis of pathway P3 that if A and C are positively correlated then 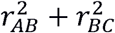 can be greater than 1. The rejection of the null model suggests that there is a relationship between FG and FI, but does not indicate whether it comes from the classical pathway or through any other source as in Fig 5c.
  g. If we ignore the non-linearity of the model and assume HOMA-IR and HOMA-β to faithfully represent insulin resistance and beta cell response, we may use the 4 predictions of the standard negative feedback pathway. It is seen that predictions from R1, R2 and R3 are accepted but the outcome of prediction from R4 is complex (Table 11c). After correcting for the effect of HOMA-IR, the FG-FI correlation should be weakened and that difference would be predicted by the correlation between HOMA-IR and FG. However instead of weakening, the FG-FI correlation becomes negative. Because of the strong positive correlation between HOMA-IR and FI, correcting for HOMA- IR subtracts from every value of FG, a quantity proportionate to FI, leading to a negative correlation between the corrected FG and FI. Additionally, simulations of the pathway show that if true insulin resistance is assumed to be correlated to FG by the same order as HOMA-IR, the correlation of true insulin resistance with FI is far less than that between HOMA-IR and FI (a result similar to Fig 6 and therefore not separately shown). Thus, there is an overfitting rejection of prediction from R1 as well. Rejection of this pathway based on two predictions is due to the unrealistically strong correlation between HOMA-IR and FI, which comes from the calculation of HOMA-IR itself.

We need to examine now to what extent HOMA-IR faithfully represents the true insulin resistance because if it does, the classical pathway certainly gets rejected. This can be examined in the simulations since the true insulin resistance is an input variable and HOMA-IR can be calculated as an outcome of the simulations. We see that HOMA-IR is correlated well with true insulin resistance when both *e*_*1*_ and *e*_*2*_ are close to zero (Fig 7). As the errors increase, the correlation becomes weaker. In the data, we do not have access to *e*_*1*_ and *e*_*2*_ but since the FG-FI correlation also becomes weaker with *e*_*2*_, we can look at how HOMA-IR represents true insulin resistance at different levels of FG-FI correlation. It can be seen that as FG-FI correlation becomes weak, HOMA-IR correlation with the true insulin resistance also becomes weak (Fig 7), but this relationship is affected by *e*_*1*_. When *e*_*1*_ is close to zero, i.e. almost all the variation in FG is explained by variation in true insulin resistance, even at low FG-FI correlation, HOMA-IR represents true insulin resistance fairly well, their correlation ranging between 0.58 and 0.7. On the other hand if we assume *e*_*1*_ to be large i.e. most of the variation in FG is due to random error or effects independent of insulin action, HOMA-IR is poorly correlated with true insulin resistance, the correlation coefficient declining to 0.2. Thus if we assume that the variance in FG is mainly caused by insulin resistance, then we have to reject the classical pathway leading to hyperinsulinemia. Alternatively, it is likely that the classical pathway is true but HOMA-IR does not represent true insulin resistance and that most of the variation in FG is not caused by insulin resistance. The substantially lower than expected slope of the FG-FI regression suggests large random errors in FG making the second interpretation more likely. In any case the classical pathway and the faithfulness of HOMA indices cannot be simultaneously true, and we have to reject at least one of them.

**Fig 7:**
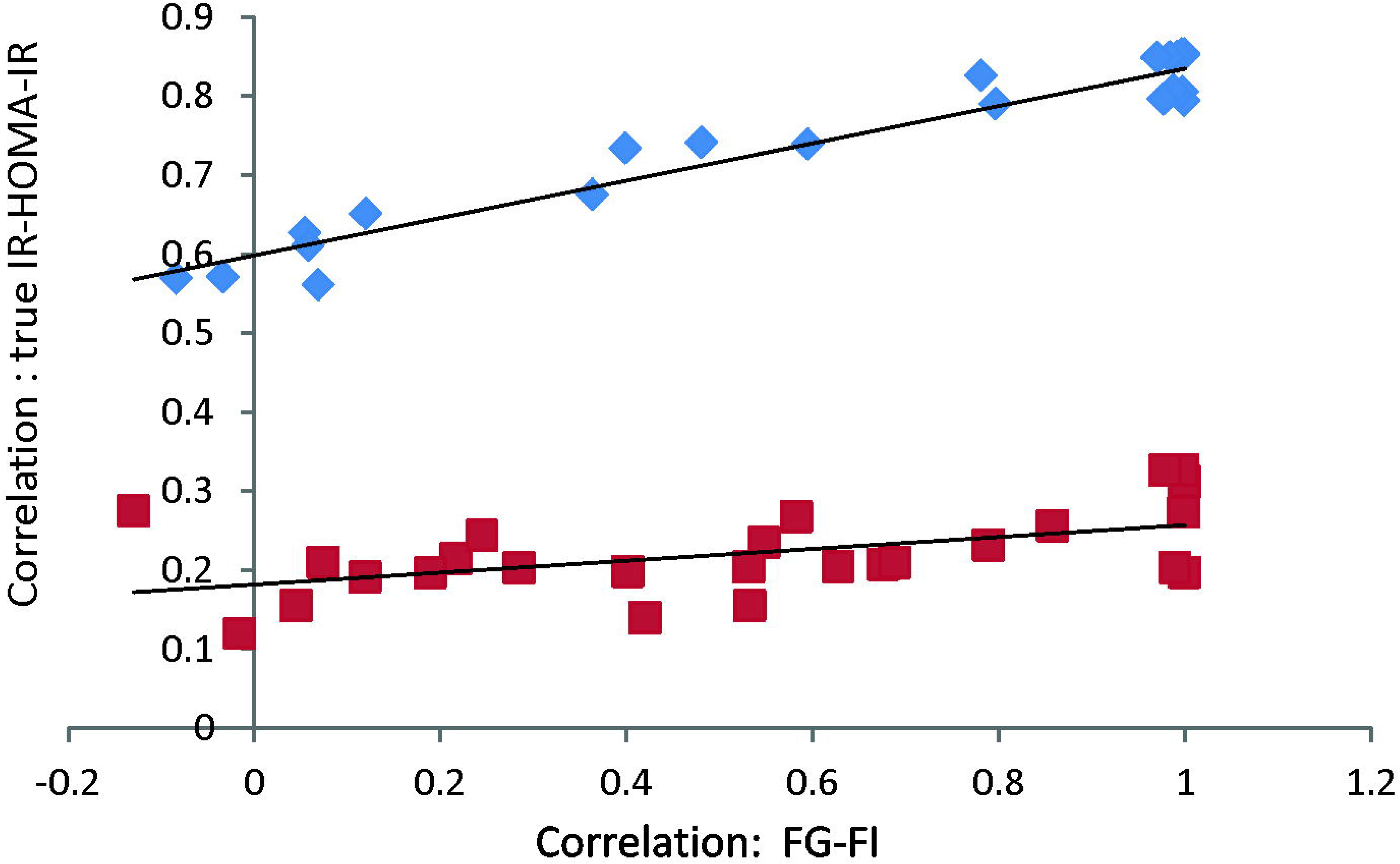
The reliability of HOMA-IR as an index of true insulin resistance: The pathway simulations were carried out at a standard deviation of *e*_*1*_=1 (blue dots) and 10 (red dots). The FG-FI correlation weakens with increase in *e*_*2*_ which also affects the correlation between true IR and HOMA-IR. It can be seen that HOMA-IR is a reliable indicator of insulin resistance when *e*_*1*_ is small, but at large *e*_*1*_ it is a poor indicator as suggested by a weak correlation with true insulin resistance.

Results of the four alternative approaches to analyze the classical pathway and the null model converge on the inference that the null model is rejected on the basis of a weak but significant correlation between FG and FI. But the weak correlation in FG and FI is not adequately explained by the classical pathway owing to multiple mismatches and rejection of many of its predictions. The pathway rejection may be partially saved by saying that HOMA-IR and HOMA- β are not good indicators of insulin resistance and beta cell response and that we do not have access to true insulin reistance to test the predictions. However the FG-FI regression slope also has a large mismatch with expectations derived from the variance in FI as well as from empirical estimates of *K*_*3*_ and *d*. Therefore, it seems more likely that FG and FI are related by causes other than the classical pathway, and HOMA-IR and HOMA- β are derived artificial constructs that do not represent any real life phenomena.

There are a number of real life interpretations of the pathway in Fig 5c. Autonomic inputs from the nervous system are known to affect both insulin secretion and liver glucose production, which might be represented by the common cause arrows of Fig 5c. Alternatively, a small error in data collection can also result in the observed FG-FI correlation. The fasting sampling is done by instructing the subjects to have no food or drink after the last evening meal. However, if even a small proportion of subjects happen to consume bed tea an hour or two before sampling, their glucose as well as insulin levels could be slightly elevated simultaneously. This can result in a weak positive correlation between FG and FI in the data. Since the fasting state is based on the honesty of the subjects and there is no independent monitoring, this source of error cannot be ignored. Thus, there are more than one possible reasons for external factors causing a weak correlation between FG and FI, and the correlation is not sufficient to support the classical pathway in the presence of multiple other mismatches.

It should be noted that the correlational patterns in the four data sets used are remarkably similar although they come from populations differing in location, ethnicity and culture. It would be important to see whether the same correlational patterns are observed in other populations as well, but we can be confident in rejecting the classical pathway at least in the populations sampled.

### What can type 2 diabetes research gain from our analysis

Putting the results together, it can safely be concluded that HOMA-IR and HOMA-β appear to be artificial constructs and reflect very marginally, if at all, the true insulin resistance and β cell response in a steady state. Until our approach for testing a causal pathway was available, there was no way to test whether HOMA-IR and HOMA-β truly represent the intended states of the system. Because of this limitation, insulin resistance was a circular argument. The inability of insulin to regulate glucose was assumed to be because of insulin resistance, but insulin resistance was measured as the inability of insulin to regulate glucose. This circularity had made the hypothesis of insulin resistance and compensatory hyperinsulinemia non-falsifiable. Our approach to pathway predictions breaks the circularity, and makes it possible to test whether the insulin resistance and glucose-mediated compensatory hyperinsulinemia hypothesis is supported by epidemiological data. At least in the populations tested, many serious anomalies in the classical pathway leading to a hyperinsulinemic, normoglycemic, insulin resistance prediabetic state are exposed. Conservatively we can argue that since HOMA-IR and HOMA-β do not represent insulin resistance and β cell response faithfully, and we do not have alternative measures for them, it may not be possible to clearly reject the classical pathway, but the data clearly show that even if true, the classical pathway has a very limited role in deciding FG, FI and their inter-relationship. Both the steady state levels have a large component of error or effects independent of the pathway under consideration. Although our analysis is restricted to the prediabetic state at present, establishing causality in the prediabetic state has implications for the over diabetic state. According to classical thinking, a failure of compensatory insulin response leads to diabetic hyperglycemia. Since our analysis questions the compensatory insulin response itself, the pathway leading to hyperglycemia is also in question. Even a highly conservative inference would demand rethinking of the causal process leading to diabetes.

Doubts about the classical pathway are raised independently by experiments using insulin receptor knockouts or insulin suppression. Muscle-specific insulin receptor knockouts show altered glucose tolerance curves but normal fasting insulin (42). Insulin suppression experiments do not result in elevated fasting glucose (43–45). Inactivation of insulin degrading enzyme raises steady state insulin levels but does not decrease glucose levels (46,47). These experiments have already challenged the classical pathway. Thus, there are multiple reasons to doubt the classical pathway. On the other hand, a number of factors other than the mutual effects of FG and FI are known to affect insulin response as well as glucose homeostasis (48–58), but these factors have not been integrated into the mainstream glucose homeostasis models. We do not intend here to test all possible alternative pathways deciding FG and FI. But our study lays down a set of methods by which this can be done, once the pathway hypotheses are clearly spelt out and the causal variables are measured. An important contribution of our methods is that physiological causal pathways can be evaluated based on epidemiological data, which is potentially a very important tool in understanding complex disorders. Experimental biology reveals what *can* happen in a system, but what *does* happen at the population level is better revealed by epidemiological data. Therefore, discerning causal signatures of pathophysiological pathways in epidemiological data is likely to be an important breakthrough.

### Conclusions

Making causal inferences from cross sectional correlational data is a long-standing problem. A correlation between two variables does not give reliable information about causal relations. However, we demonstrate here, in the context of steady state homeostatic systems, using mathematical proofs as well as simulations from causal pathways that, in a set of three or more correlated variables, it is possible to test causal hypotheses based on the interrelationships of regression-correlation parameters. This is potentially a highly valuable tool in making causal inferences from cross sectional data in several fields.

Using this set of principles, we tested the classical causal assumption behind the hyperinsulinemic, normoglycemic, insulin resistant or pre-diabetic state. The analysis showed that this causal pathway and the measures of insulin resistance and insulin response were not supported by epidemiological data. Thus, the objections raised recently to the classical causal pathway are validated and alternative causal pathways that already have substantial experimental evidence need to be integrated in the mainstream clinical thinking.

## Acknowledgements

We thank Chittaranjan Yajnik for making the CRISIS and PMNS data sets available to us. We would also like to thank Louise Hayes and Raj Bhopal for making the data from NHP studies available. We thank Subhash Lele and Anil Gore for useful comments on an earlier draft manuscript. We also thank Rajiv Gandhi Science and Technology Commission, (RGSTC), Maharashtra State, India for partial support.

## Supporting Information

**S1 Text.** Deriving predictions for regression parameters based on causal equations and calculation of causal parameters for simulations.

**S2 Text.** Empirical estimates for parameters used in simulations.

## References

1. Baker JP. Mercury, Vaccines, and Autism. Am J Public Health [Internet]. 2008 |mFeb;98(2):244–53. Available from: http://ajph.aphapublications.org/doi/10.2105/AJPH.2007.113159

2. DeStefano F. Vaccines and Autism: Evidence Does Not Support a Causal Association. Clin Pharmacol Ther [Internet]. 2007 Dec 10;82(6):756–9. Available from: http://doi.wiley.com/10.1038/sj.clpt.6100407

3. Parascandola M, Weed DL. Causation in epidemiology. J Epidemiol Community Health. 2001;55(12):905–12.

4. Gerber JS, Offit PA. Vaccines and Autism: A Tale of Shifting Hypotheses. Clin Infect Dis [Internet]. 2009 Feb 15;48(4):456–61. Available from: https://academic.oup.com/cid/article-lookup/doi/10.1086/596476

5. Ratzan SC. Setting the Record Straight: Vaccines, Autism, and the Lancet. J Health Commun [Internet]. 2010 Apr 30;15(3):237–9. Available from: http://www.tandfonline.com/doi/abs/10.1080/10810731003780714

6. Ejima K, Li P, Smith DL, Nagy TR, Kadish I, van Groen T, et al. Observational research rigour alone does not justify causal inference. Eur J Clin Invest [Internet]. 2016 Dec;46(12):985–93. Available from: http://doi.wiley.com/10.1111/eci.12681

7. Hill AB. The Environment and Disease: Association or Causation? Proc R Soc Med. 1965 May;58(5):295–300.

8. Meehl PE, Waller NG. The path analysis controversy: a new statistical approach to strong appraisal of verisimilitude. Psychol Methods [Internet]. 2002 Sep;7(3):283–300. Available from: http://www.ncbi.nlm.nih.gov/pubmed/12243300

9. Niles HE. The Method of Path Coefficients an Answer to Wright. Genetics [Internet]. 1923 May;8(3):256–60. Available from: http://www.ncbi.nlm.nih.gov/pubmed/17246012

10. Wright S. The Method of Path Coefficients. Ann Math Stat [Internet]. 1934 Sep;5(3):161–215. Available from: http://projecteuclid.org/euclid.aoms/1177732676

11. Wright S. Path Coefficients and Path Regressions: Alternative or Complementary Concepts? Biometrics [Internet]. 1960 Jun;16(2):189. Available from: http://www.jstor.org/stable/2527551?origin=crossref

12. Greenland S. An introduction to instrumental variables for epidemiologists. Int J Epidemiol [Internet]. 2000 Aug;29(4):722–9. Available from: https://academic.oup.com/ije/article-lookup/doi/10.1093/ije/29.4.722

13. Granger CWJ. Investigating Causal Relations by Econometric Models and Cross-spectral Methods. Econometrica [Internet]. 1969 Aug;37(3):424. Available from: http://www.jstor.org/stable/1912791?origin=crossref

14. Rosenbaum PR, Rubin DB. The central role of the propensity score in observational studies for causal effects. Biometrika [Internet]. 1983;70(1):41–55. Available from: https://academic.oup.com/biomet/article-lookup/doi/10.1093/biomet/70.1.41

15. Peters J, Janzing D, Scholkopf B. Causal Inference on Discrete Data Using Additive Noise Models. IEEE Trans Pattern Anal Mach Intell. 2011 Dec;33(12):2436–50.

16. Neuberg LG. CAUSALITY: MODELS, REASONING, AND INFERENCE, by Judea Pearl, Cambridge University Press, 2000. Econom Theory [Internet]. 2003 Aug 6;19(4). Available from: http://www.journals.cambridge.org/abstract_S0266466603004109

17. Pearl J. Causal inference in statistics: An overview. Stat Surv [Internet]. 2009;3:96–146. Available from: http://projecteuclid.org/euclid.ssu/1255440554

18. Peters J, Janzing D, Scholkopf B. Causal Inference on Discrete Data Using Additive Noise Models. IEEE Trans Pattern Anal Mach Intell [Internet]. 2011 Dec;33(12):2436–50. Available from: http://ieeexplore.ieee.org/document/5740928/

19. Peters J, Mooij JM, Janzing D, Schölkopf B. Causal Discovery with Continuous Additive Noise Models. J Mach Learn Res. 2014;15:2009–53.

20. Eichler M. Causal inference with multiple time series: principles and problems. Philos Trans R Soc A Math Phys Eng Sci [Internet]. 2013 Jul 15;371(1997):20110613–20110613. Available from: http://rsta.royalsocietypublishing.org/cgi/doi/10.1098/rsta.2011.0613

21. Abdul-Ghani MA, Tripathy D, DeFronzo RA. Contributions of beta-cell dysfunction and insulin resistance to the pathogenesis of impaired glucose tolerance and impaired fasting glucose. Diabetes Care [Internet]. 2006 May;29(5):1130–9. Available from: http://www.ncbi.nlm.nih.gov/pubmed/16644654

22. Cerf ME. Beta Cell Dysfunction and Insulin Resistance. Front Endocrinol (Lausanne)[Internet]. 2013;4. Available from: http://journal.frontiersin.org/article/10.3389/fendo.2013.00037/abstract

23. Watve M. Doves, Diplomats, and Diabetes [Internet]. New York, NY: Springer New York; 2013. Available from: http://link.springer.com/10.1007/978-1-4614-4409-1

24. Dubuc PU. The development of obesity, hyperinsulinemia, and hyperglycemia in ob/ob mice. Metabolism [Internet]. 1976 Dec;25(12):1567–74. Available from: http://www.ncbi.nlm.nih.gov/pubmed/994838

25. Dubuc PU. Non-essential role of dietary factors in the development of diabetes in ob/ob mice. J Nutr [Internet]. 1981 Oct;111(10):1742–8. Available from: http://www.ncbi.nlm.nih.gov/pubmed/7026742

26. Garvey WT, Olefsky JM, Marshall S. Insulin induces progressive insulin resistance in cultured rat adipocytes. Sequential effects at receptor and multiple postreceptor sites. Diabetes [Internet]. 1986 Mar;35(3):258–67. Available from: http://www.ncbi.nlm.nih.gov/pubmed/3512337

27. Shanik MH, Xu Y, Skrha J, Dankner R, Zick Y, Roth J. Insulin Resistance and Hyperinsulinemia: Is hyperinsulinemia the cart or the horse? Diabetes Care [Internet].2008 Feb 1;31(Supplement 2):S262–8. Available from:http://care.diabetesjournals.org/cgi/doi/10.2337/dc08-s264

28. Corkey BE. Banting lecture 2011: hyperinsulinemia: cause or consequence? Diabetes [Internet]. 2012 Jan;61(1):4–13. Available from: http://www.ncbi.nlm.nih.gov/pubmed/22187369

29. Lerner RL, Porte D. Acute and steady-state insulin responses to glucose in nonobese diabetic subjects. J Clin Invest [Internet]. 1972 Jul 1;51(7):1624–31. Available from: http://www.jci.org/articles/view/106963

30. Turner RC, Holman RR, Matthews D, Hockaday TD, Peto J. Insulin deficiency and insulin resistance interaction in diabetes: estimation of their relative contribution by feedback analysis from basal plasma insulin and glucose concentrations. Metabolism [Internet]. 1979 Nov;28(11):1086–96. Available from: http://www.ncbi.nlm.nih.gov/pubmed/386029

31. Halter JB, Ward WK, Porte D, Best JD, Pfeifer MA. Glucose regulation in non-insulin-dependent diabetes mellitus. Interaction between pancreatic islets and the liver. Am J Med [Internet]. 1985 Aug 23;79(2B):6–12. Available from: http://www.ncbi.nlm.nih.gov/pubmed/2863979

32. Matthews DR, Hosker JP, Rudenski AS, Naylor BA, Treacher DF, Turner RC. Homeostasis model assessment: insulin resistance andLJ?-cell function from fasting plasma glucose and insulin concentrations in man. Diabetologia [Internet]. 1985 Jul;28(7):412–9. Available from: http://link.springer.com/10.1007/BF00280883

33. WA Fuller, editor. Measurement Error Models [Internet]. Hoboken, NJ, USA: John Wiley & Sons, Inc.; 1987. (Wiley Series in Probability and Statistics). Available from: http://doi.wiley.com/10.1002/9780470316665

34. Griggs RC, Kingston W, Jozefowicz RF, Herr BE, Forbes G, Halliday D. Effect of testosterone on muscle mass and muscle protein synthesis. J Appl Physiol [Internet]. 1989 Jan;66(1):498–503. Available from: http://www.ncbi.nlm.nih.gov/pubmed/2917954

35. Kraemer WJ, Ratamess NA. Hormonal responses and adaptations to resistance exerciseand training. Sports Med [Internet]. 2005;35(4):339–61. Available from:http://www.ncbi.nlm.nih.gov/pubmed/15831061

36. Cumming DC, Brunsting LA, Strich G, Ries AL, Rebar RW. Reproductive hormone increases in response to acute exercise in men. Med Sci Sports Exerc [Internet]. 1986 Aug;18(4):369–73. Available from: http://www.ncbi.nlm.nih.gov/pubmed/2943968

37. Tomasi T, Sledz D, Wales JK, Recant L. Insulin half-life in normal and diabetic subjects. Rev Neuropsychiatr Infant [Internet]. 1966 Dec;14(12):315–7. Available from: http://www.ncbi.nlm.nih.gov/pubmed/5988016

38. Pories WJ, Dohm GL. Diabetes: Have We Got It All Wrong?: Hyperinsulinism as the culprit: surgery provides the evidence. Diabetes Care [Internet]. 2012 Dec 1;35(12):2438–42. Available from: http://care.diabetesjournals.org/cgi/doi/10.2337/dc12-0684

39. Yajnik CS, Joglekar C V, Lubree HG, Rege SS, Naik SS, Bhat DS, et al. Adiposity,inflammation and hyperglycaemia in rural and urban Indian men: Coronary Risk of Insulin Sensitivity in Indian Subjects (CRISIS) Study. Diabetologia [Internet]. 2008 Jan;51(1):39–46. Available from: http://www.ncbi.nlm.nih.gov/pubmed/17972060

40. Bhopal R, Unwin N, White M, Yallop J, Walker L, KGMM Alberti, et al. Heterogeneityof coronary heart disease risk factors in Indian, Pakistani, Bangladeshi, and European origin populations: cross sectional study. BMJ [Internet]. 1999 Jul 24;319(7204):215–20.Available from: http://www.bmj.com/cgi/doi/10.1136/bmj.319.7204.215

41. Joshi NP, Kulkarni SR, Yajnik CS, Joglekar C V., Rao S, Coyaji KJ, et al. Increasing maternal parity predicts neonatal adiposity: Pune Maternal Nutrition Study. Am J Obstet Gynecol. 2005;193(3):783–9.

42. Kadowaki T. Insights into insulin resistance and type 2 diabetes from knockout mouse models. J Clin Invest [Internet]. 2000 Aug 15;106(4):459–65. Available from: http://www.jci.org/articles/view/10830

43. Gill G V., Rauf O, MacFarlane IA. Diazoxide treatment for insulinoma: a national UK survey. Postgrad Med J [Internet]. 1997 Oct 1;73(864):640–1. Available from: http://pmj.bmj.com/cgi/doi/10.1136/pgmj.73.864.640

44. Alemzadeh R, Karlstad MD, Tushaus K, Buchholz M. Diazoxide enhances basal metabolic rate and fat oxidation in obese Zucker rats. Metabolism [Internet]. 2008 Nov;57(11):1597–607. Available from: http://linkinghub.elsevier.com/retrieve/pii/S0026049508002382

45. Alemzadeh R, Tushaus KM. Modulation of Adipoinsular Axis in Prediabetic Zucker Diabetic Fatty Rats by Diazoxide. Endocrinology [Internet]. 2004 Dec;145(12):5476–84. Available from: https://academic.oup.com/endo/article-lookup/doi/10.1210/en.2003-1523

46. Maianti JP, McFedries A, Foda ZH, Kleiner RE, Du XQ, Leissring MA, et al. Anti-diabetic activity of insulin-degrading enzyme inhibitors mediated by multiple hormones. Nature [Internet]. 2014 Jul 21;511(7507):94–8. Available from:http://www.nature.com/articles/nature13297

47. Abdul-Hay SO, Kang D, McBride M, Li L, Zhao J, Leissring MA. Deletion of insulin-degrading enzyme elicits antipodal, age-dependent effects on glucose and insulin tolerance. PLoS One [Internet]. 2011;6(6):e20818. Available from: http://www.ncbi.nlm.nih.gov/pubmed/21695259

48. Levinger I, Goodman C, Matthews V, Hare DL, Jerums G, Garnham A, et al. BDNF, Metabolic Risk Factors, and Resistance Training in Middle-Aged Individuals. Med Sci Sport Exerc [Internet]. 2008 Mar;40(3):535–41. Available from: https://insights.ovid.com/crossref?an=00005768-200803000-00020

49. Reinehr T, Roth CL. A new link between skeleton, obesity and insulin resistance: relationships between osteocalcin, leptin and insulin resistance in obese children before and after weight loss. Int J Obes [Internet]. 2010 May 12;34(5):852–8. Available from: http://www.nature.com/articles/ijo2009282

50. Polgreen LE, Jacobs DR, Nathan BM, Steinberger J, Moran A, Sinaiko AR. Association of Osteocalcin With Obesity, Insulin Resistance, and Cardiovascular Risk Factors in Young Adults. Obesity [Internet]. 2012 Nov;20(11):2194–201. Available from: http://doi.wiley.com/10.1038/oby.2012.108

51. Lo C-M, Obici S, Dong HH, Haas M, Lou D, Kim DH, et al. Impaired insulin secretionand enhanced insulin sensitivity in cholecystokinin-deficient mice. Diabetes [Internet]. 2011 Jul;60(7):2000–7. Available from: http://www.ncbi.nlm.nih.gov/pubmed/21602512

52. Han DH, Hansen PA, Chen MM, Holloszy JO. DHEA treatment reduces fat accumulation and protects against insulin resistance in male rats. J Gerontol A Biol Sci Med Sci[Internet]. 1998 Jan;53(1):B19-24. Available from: http://www.ncbi.nlm.nih.gov/pubmed/9467418

53. Pitteloud N, Mootha VK, Dwyer AA, Hardin M, Lee H, Eriksson K-F, et al. Relationship between testosterone levels, insulin sensitivity, and mitochondrial function in men. Diabetes Care [Internet]. 2005 Jul;28(7):1636–42. Available from: http://www.ncbi.nlm.nih.gov/pubmed/15983313

54. Polderman KH, Gooren LJ, Asscheman H, Bakker A, Heine RJ. Induction of insulin resistance by androgens and estrogens. J Clin Endocrinol Metab [Internet]. 1994 Jul;79(1):265–71. Available from: http://www.ncbi.nlm.nih.gov/pubmed/8027240

55. Nonogaki K. New insights into sympathetic regulation of glucose and fat metabolism. Diabetologia [Internet]. 2000 May;43(5):533–49. Available from: http://www.ncbi.nlm.nih.gov/pubmed/10855527

56. Meek TH, Wisse BE, Thaler JP, Guyenet SJ, Matsen ME, Fischer JD, et al. BDNF action in the brain attenuates diabetic hyperglycemia via insulin-independent inhibition of hepatic glucose production. Diabetes [Internet]. 2013 May;62(5):1512–8. Available from: http://www.ncbi.nlm.nih.gov/pubmed/23274899

57. Holmaeng A, Bjoerntorp P. The effects of testosterone on insulin sensitivity in male rats. Acta Physiol Scand [Internet]. 1992 Dec;146(4):505–10. Available from: http://doi.wiley.com/10.1111/j.1748-1716.1992.tb09452.x

58. Parton LE, Ye CP, Coppari R, Enriori PJ, Choi B, Zhang C-Y, et al. Glucose sensing by POMC neurons regulates glucose homeostasis and is impaired in obesity. Nature [Internet]. 2007 Sep 29;449(7159):228–32. Available from: http://www.nature.com/articles/nature06098

